# Using nanopore sequencing to obtain complete genomes from saliva samples

**DOI:** 10.1101/2022.05.19.492709

**Authors:** Jonathon L. Baker

## Abstract

Obtaining complete, high-quality reference genomes is essential to the study of any organism. Recent advances in nanopore sequencing, as well as genome assembly and analysis methods, have made it possible to obtain complete bacterial genomes from metagenomic (i.e. multispecies) samples, including those from the human microbiome. In this study, methods are presented to obtain complete bacterial genomes from human saliva using complementary Oxford Nanopore (ONT) and Illumina sequencing. Applied to 3 human saliva samples, these methods resulted in 11 complete bacterial genomes: 3 *Saccharibacteria* clade G6 (aka Ca. Nanogingivalaceae HMT-870), 1 *Saccharibacteria* clade G1 HMT-348, 2 *Rothia mucilaginosa*, 2 *Actinomyces graevenitzii*, 1 *Mogibacterium diversum*, 1 *Lachnospiraceae* HMT-096, 1 *Lancefieldella parvula*, and one circular chromosome of Ruminococcaceae HMT-075 (which likely has at least 2 chromosomes). The 4 *Saccharibacteria* genomes, as well as the *Actinomyces graeventizii* genomes, represented the first complete genomes from their respective bacterial taxa. Aside from the complete genomes, the assemblies contained 147 contigs of over 500,000 bp each and thousands of smaller contigs, together representing many additional draft genomes including many which are likely near-complete. The complete genomes enabled highly accurate pangenome analysis, which identified unique and missing features of each genome compared to its closest relatives with complete genomes available in public repositories. These features provide clues as to the lifestyle and ecological role of these bacteria within the human oral microbiota, which will be particularly useful in designing future studies of the specific taxa that have never been isolated or cultivated.

## Introduction

Sequencing the genomes of the taxa comprising the human microbiome is fundamental to our understanding of how these species live and affect our health (Gill et al., 2006; Human Microbiome Project, 2012). For a given taxon, obtaining a high-quality reference genome is crucial for several reasons. First, a high-quality genome allows researchers to quantify the abundance of a particular taxon, or its mRNA transcripts, in microbiome samples accurately through read-mapping (Venter et al., 2004). Second, with a complete genome in hand, researchers can predict the metabolic and ecological role of the taxon, which can be especially important for species which have not yet been isolated or cultivated in the lab (which still represents the majority of bacteria) (Fleischmann et al., 1995). Finally, a high-quality genome is critical to guide wet lab experimentation, such as genome editing and mutagenesis (Fleischmann et al., 1995). Despite the vast and continuously growing number of microbial genomes in databases such as NCBI RefSeq and the Human Oral Microbiome Database (HOMD), many bacteria, including those of the human oral microbiome, lack reference genomes. Complete genomes, in the case of bacteria, are most frequently circular chromosomes, although some taxa have linear chromosomes, and some taxa, such as *Prevotella spp*., have more than one chromosome (Naito et al., 2016).

For the past ∼15 years, Illumina shotgun sequencing has been the industry workhorse, revolutionizing the life sciences by lowering the cost and increasing the throughput of sequencing by several orders of magnitude, while providing very high accuracy (Bennett, 2004; Bentley et al., 2008). The main drawback to this technology is the short length of the reads, which are generally 150 or 300 bp. To obtain genomes or metagenomes (i.e. genomic sequencing of samples congaing the genomes of multiple taxa or isolates, such as a microbiome sample), these short reads must be assembled. However, most genomes have repetitive regions such as ribosomal RNA regions or invertases, that can be well over 10,000 bp in length (and may be contiguous or in distant loci, both are problematic). The short-reads from within these regions all map non-specifically, therefore one cannot tell how many repeats exist within the repeat structure, or what part of the genome connects on the other side of the repeat (Figure 1). Consequently, these regions cannot be elucidated with confidence during assembly, causing the production of separate fragments, known as contigs, rather than a complete chromosome in the resulting draft genome (Athanasopoulou, Boti, Adamopoulos, Skourou, & Scorilas, 2021). When dealing with metagenomes, rather than isolate genomes, several additional problems are introduced (Figure 1). First, as there are a finite number of reads generated in a sequencing run, more genomes in the sample will mean fewer reads produced per genome (Venter et al., 2004). Because accurate genome assembly is dependent on coverage depth, fewer reads supporting a given genome will lead to a more fragmented, incomplete, and error prone assembly. Second, it can be difficult to know which particular contigs are from the same genome. “Binning” is the process to sort the metagenomic contigs into discrete, fragmented draft genomes, and many computational tools have been developed to do this, frequently using k-mer frequency, coverage depth, GC content, and/or alignment to references (Alneberg et al., 2014; Kang et al., 2019; Wu, Simmons, & Singer, 2016). However, even the most exhaustive automated and manual binning strategies can “misplace” contigs into the wrong genome bin, leading to “contamination” (L. X. Chen, Anantharaman, Shaiber, Eren, & Banfield, 2020). It can be especially problematic when these contaminated draft genomes make it as far as public repositories, as downstream researchers will usually accept these assemblies as gospel and use the erroneous data to design further experiments (Shaiber & Eren, 2019). Complete genomes (i.e. contiguous chromosomes) are, by definition, free of these types of errors, and are therefore the most useful tools for scientists, as high confidence can be placed in their accuracy. It is imperative that researchers examine whether the reference genomes in-use are ‘complete’ or ‘draft’, and recognize the limitations of draft genomes (L. X. Chen et al., 2020; Shaiber & Eren, 2019).

**Figure 1:**
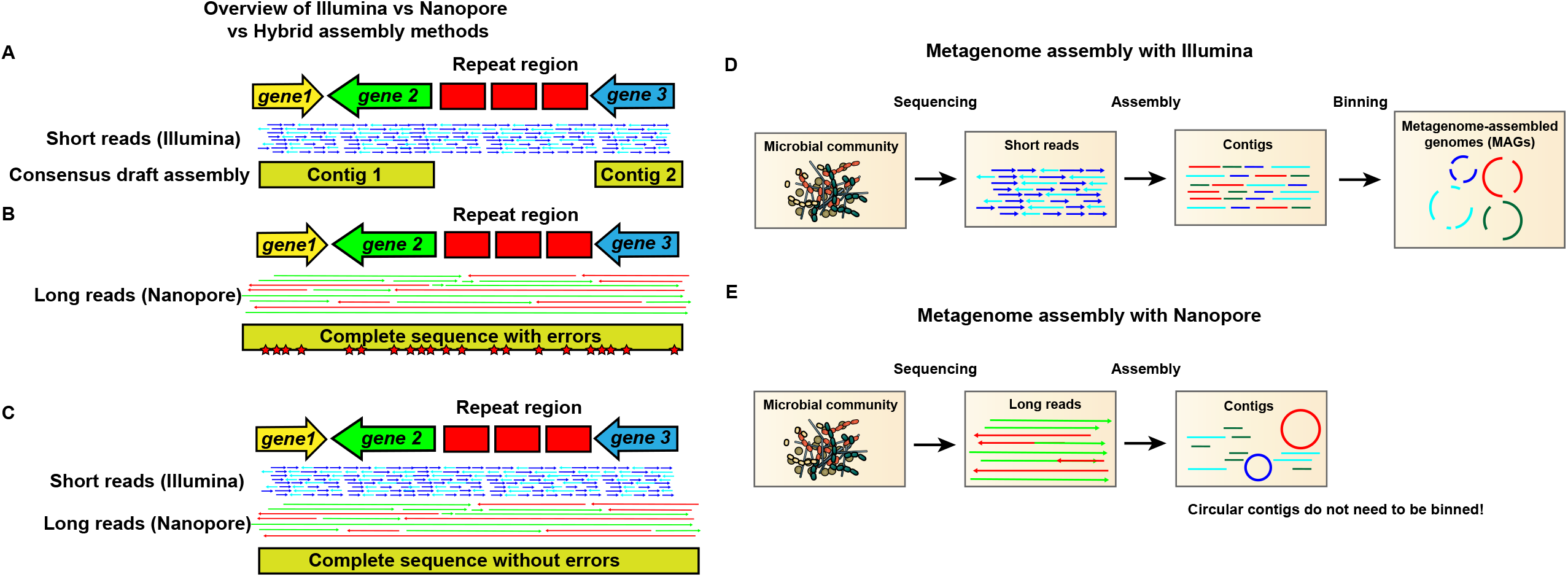
Getting complete and accurate genomes from metagenomes using Illumina and Nanopore sequencing. Illustrations showing assembly of genomes using (A) Illumina only, (B) Nanopore only, or (C) both Illumina and Nanopore. (A) Assembly with short reads yields high accuracy, but short-reads map nonspecifically in repeat regions, which therefore cannot be resolved. The result is an assembly fragmented into contigs. (B) Assembly with nanopore reads resolves repeat regions, because individual reads map all the way through, resulting in a complete, contiguous assembly. Errors will be present due to the higher error rate of nanopore, most frequently in homopolymeric tracts and short repeats. (C) Assembly with complementary Illumina and nanopore sequencing allows for nanopore errors to be corrected using the more accurate short-reads, yielding an assembly that is complete, contiguous, and highly accurate. (D). Metagenome assembly with Illumina sequencing requires binning that frequently produces contamination in the resulting fragmented draft metagenome-assembled genome. (E) Metagenome assembly with ONT produces circular contigs that represent complete chromosomes, and therefore do not need to be binned.

Until recently, due to the constraints inherent in short-read sequencing just described, obtaining complete genomes usually required [1] manual steps beyond computational assembly, such as PCR and further sequencing, due to the repeat regions mentioned above, and [2] pure cultures to eliminate the problems inherent with sequencing metagenomes. These requirements significantly limited the output of complete genomes, particularly since the majority of bacterial species have yet to be successfully isolated and cultured in the lab. However, recent advancements in long-read sequencing technology, particularly that of Oxford Nanopore Technologies (ONT), have been revolutionary (Jain et al., 2015). Unlike the sequencing-by-synthesis techniques employed by Illumina and PacBio technologies, ONT sequences native DNA or RNA molecules, and read length is only limited by the size of input DNA, allowing for read lengths of over 1 Mbp under the right conditions (Jain et al., 2015). The long reads produced by ONT sequencing easily map all the way through problematic repeat regions, significantly improving assembly contiguity and frequently resulting in complete, circular genomes/chromosomes, even from metagenomic (i.e. non-isolate) samples, eliminating the need for binning (Figure 1) (Cusco, Perez, Vines, Fabregas, & Francino, 2021; Loman, Quick, & Simpson, 2015; Moss, Maghini, & Bhatt, 2020). Recently, ONT sequencing was instrumental in obtaining the first telomere-to-telomere complete human genome (Nurk et al., 2022).

In ONT sequencing, native nucleic acids are ratcheted through a nanopore embedded in a synthetic membrane, and the changes in electrical current are monitored (Jain et al., 2015). Sequences of bases or windows of bases cause specific and predictable perturbations in the electrical potential, therefore the sequence of bases passing through the nanopore can be reconstructed by analyzing the current (Amarasinghe et al., 2020; Athanasopoulou et al., 2021). Although nanopore sequencing technology is not new, its applicability was limited for many years by a high error rate. These error rates have dropped substantially in recent years, from 35-40% in 2015 to <1% with current ONT instrumentation and software (Liu et al., 2021). These improvements have been due to major advancements in both nanopore chemistry and in machine learning technology, which underpins the basecalling algorithms. The errors introduced by nanopore sequencing are not random, but typically occur during homopolymeric tracts or short repeats, where the basecalling software has difficulty identifying how many consecutive iterations of a given base or bases have passed through the nanopore (Amarasinghe et al., 2020). This leads to insertions or deletions (i.e. indels) in these homopolymeric tracts, which can cause apparent frameshifts, and therefore can have a significant impact on downstream gene calling and gene annotation (Amarasinghe et al., 2020). Increasing depth of sequencing coverage mitigates, but does not eliminate this issue. Consequently, at this time ONT sequencing can fully stand on its own for many applications, such as RNA-seq, and assembly of draft genomes with a high degree of completeness (Athanasopoulou et al., 2021). ONT sequencing is also useful to perform 16S amplicon sequencing to profile microbial communities, as the longer ONT reads can span the entire 16 rRNA gene, therefore providing coverage across all the variable regions, which allows for increased taxonomic specificity and less taxonomic bias compared to methods targeting only one variable region (Matsuo et al., 2021). However, for producing error free, publication quality complete genomes, complementary Illumina sequencing is still helpful, as the substantially lower error rate in Illumina reads can be used to correct errors in the ONT-based assembly (Figure 1) (Athanasopoulou et al., 2021).

In this study, a protocol for generating complete genomes of oral bacteria from saliva using complementary ONT and Illumina sequencing is reported. Sequencing of 3 saliva samples resulted in 12 complete, circular chromosomes. Among these were 3 genomes, representing two distinct species, of clade G6 *Saccharibacteria* (HMT-870) (Baker, 2021a, 2021b), 1 genome of clade G1 *Saccharibacteria* HMT-348 (Baker, 2022), 2 genomes of *Actinomyces graevenitzii*, 2 genomes of *Rothia mucilaginosa*, 1 genome of *Lachnospiraceae* HMT-096, 1 genome of *Mogibacterium diversum*, 1 genome of *Lancefieldella parvula*, and also a complete circular chromosome from Ruminococcaceae HMT-075 (this taxon likely has multiple chromosomes). The properties of these genomes are then highlighted.

## Results and Discusssion

### Extracting high molecular weight genomic DNA (HMW gDNA)

During ONT sequencing, the native DNA or RNA molecules are sequenced, and the read length of the sequences produced is limited only by the length of the input material. Therefore, to obtain the most complete genomes possible from genomic or metagenomic sequencing, it is important to obtain HMW gDNA with as little shearing of the molecules as possible. For the Ligation Sequencing Kit used to prepare the DNA library for ONT sequencing, 1 μg of HMW gDNA is required. Several gDNA preparation methods were tested here, described in more detail in the Materials and Methods section. The protocol which gave the best results, and was used subsequently here, was a phenol:chloroform-based protocol originally published by Chen and Burne (Y. Y. Chen, Clancy, & Burne, 1996), which was recently optimized specifically for nanopore sequencing of *Streptococcus mutans* B04Sm5 (Baker & Edlund, 2020). It should be emphasized that during the course of this study several new HMW isolation kits/protocols have been released that may produce even longer reads (and therefore additional complete genomes) but were not tested here. These include the Ultra-Long DNA Sequencing Kit from Oxford Nanopore Technologies, Inc. and the Nanobind UHMW DNA Extraction protocol from Circulomics, Inc.

The 3 saliva samples sequenced with ONT in this study were SC23, SC24, and SC33, which were all collected from children with healthy dentition in Los Angeles, CA as previously described (NCBI accession number PRJNA624185) (Baker, 2021a, 2021b). These samples were previously sequenced using Illumina technology and subjected to metagenomic analysis (Baker et al., 2021). The corresponding Illumina sequencing libraries from these samples are available in the Sequence Read Archive (SRA) with accession numbers SRX4318838 (SC23), SRX4318837 (SC24), and SRX4318835 (SC33). Here, HMW gDNA was extracted from 1 ml aliquots of the same saliva samples used to produce the original Illumina short-read libraries using the modified Chen and Burne protocol (Y. Y. Chen et al., 1996) described above and in further detail in the Materials and Methods. The samples were sequenced on a GridION instrument with each sample using a full R9.4.1 flow cell. ONT sequencing of SC23 yielded 3.2 million reads with an N50 of 13,719 bp, while nanopore sequencing of SC24 yielded 3.9 million reads with an N50 of 14,073 bp, and nanopore sequencing of SC33 yielded 8.9 million reads with an N50 of 6,527 bp. Because the saliva contains human DNA, which will only complicate the assembly process for the microbial genomes, human reads were removed from all 3 long-read libraries by mapping the read libraries to the human genome using minimap2 (Li, 2018) and removing the reads that mapped. Following removal of the human reads, SC23 contained 686K reads with an N50 of 19,172 bp, SC24 contained 1.07M reads with an N50 of 19,058 bp, and SC33 contained 1.2M reads with an N50 of 13,393 bp. The longest single reads in SC23, SC24, and SC33 were 88,225 bp, 109,498 bp, and 124,167 bp, respectively.

### Metagenomic assembly

Each of the 3 long-read libraries was assembled using meta-Flye (Kolmogorov et al., 2020). Tables S1-S3 contain a summary of the assembly info from Flye. In terms of putative complete genomes, considered here to be a circular contig >500 Kbp, SC23 had 2, SC24 had 5, and SC33 had 4. These assemblies are described in Table 1. The assemblies also contained many very large linear contigs (>500,000); SC23 had 38, SC24 had 69, and SC33 had 40, with the largest linear contigs in all 3 assemblies being well over 2 Mbp; likely representing near-complete genomes (or possibly complete genomes from organisms which have linear, rather than circular, genomes). As the circular contigs are much more likely to be complete genomes (although not error-free at this stage), only large circular contigs are the focus of this study.

**Table 1:**
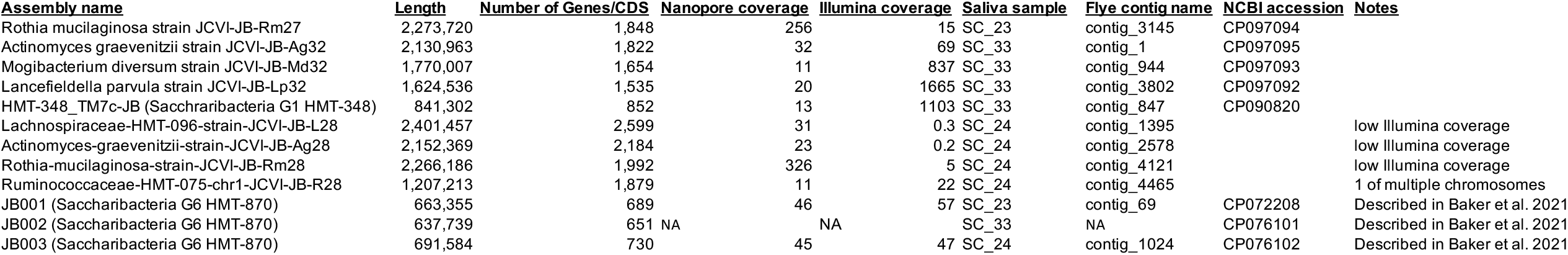
Genomes of interest produced by this study.

### Assembly of JB001, JB002, and JB003 (*Saccharibacteria* clade G6)

Candidatus Nanogingivalaceae (a Clade G6 *Saccharibacteria* / HMT-870) strains JB001, JB002, and JB003 were derived from this dataset and have already been described previously (Baker, 2021a, 2021b), however it is useful to summarize the assembly methods used on those genomes to explain how the assembly pipeline of the genomes described in this study matured and improved over the course of the study (Figure 2A). Assembly methods using both long and short reads either [1] assemble the long reads into a draft and then remove errors in the long reads using the short reads or [2] assemble the short reads and then use long reads to bridge together the disjointed contigs. Assembly with Flye, followed by short-read polishing uses the former strategy, while assembling both long and short reads mapping to the draft assembly using Unicycler (Wick, Judd, Gorrie, & Holt, 2017), employs the latter. Both of these methods were attempted, and then Trycycler (Wick et al., 2021) was used to determine the best final consensus assembly using the 2 draft assemblies. The Trycycler consensus assembly was then polished with the long reads using Medaka (https://github.com/nanoporetech/medaka) and with the short reads using Pilon (Walker et al., 2014) (Figure 2A).

**Figure 2:**
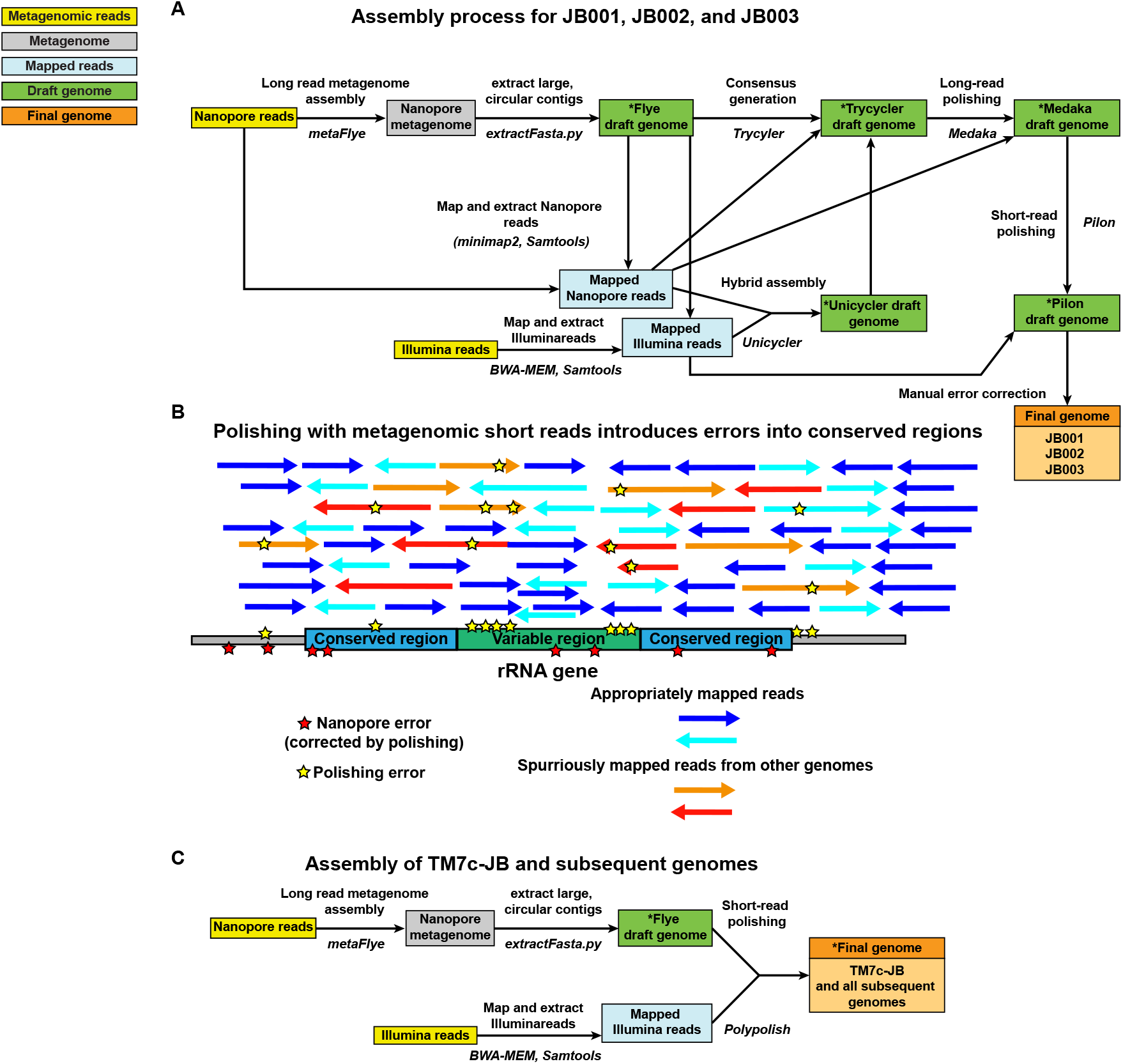
Genome assembly and polishing pipelines used in this study. (A) Original pipeline of assembling and correcting errors in the metagenome-assembled genomes JB001, JB002, and JB003. (B) Polishing with metagenomic short reads introduces errors into conserved regions. This issue was identified with the pipeline presented in Panel A above. Metagenomic short reads from other species/genomes spuriously map to the draft genome because of conserved regions, such as those found in rRNA. These spuriously mapped reads introduce errors into the polished genome. The polishing is still useful for correcting nanopore errors elsewhere in the genome, however. (C) Pipeline used on HMT-348-TM7c-JB and all other genomes in this study. Polypolish alleviates the issue described in Panel B almost entirely.

However, comparison of annotated versions of the draft genomes both before and after polishing steps, along with reference rRNA genes from Candidatus Nanogingivalaceae, revealed a problem (Figure 2B) (Baker, 2021b). While genomic rRNA sequences from the original Flye assembly were nearly identical to the HOMD reference rRNA sequences, the genomic rRNA sequences produced following polishing steps contained many more mismatches (Baker, 2021b). The likely explanation is that because regions of the rRNA are highly conserved (which is why 16S amplicon sequencing is a widely used and cost-effective means of microbiome analysis), and the read libraries used for Medaka and Pilon polishing came from a metagenomic sample, reads from other species/genomes align to the conserved rRNA regions and get spuriously mapped to the draft assembly (Baker, 2021b). The portions of the reads that do not perfectly align the assembly then cause errors to be introduced during the polishing (Figure 2B). Outside of the widely conserved rRNA operons, however, the Medaka and Pilon polishing steps were useful in correctly removing errors, especially the frameshift-causing indels within homopolymeric tracts, described above as a drawback to ONT sequencing (Baker, 2021b). To solve this issue, the sequence of the rRNA regions within the consensus genome was manually changed back to that of the original, correct Flye assembly, while the remainder of the genome kept the changes introduced by polishing (Baker, 2021b). This protocol was used in the production of JB001, JB002, and JB003 (all Candidatus Nanogingivalacae HMT-870) (Baker, 2021a, 2021b).

### Assembly of HMT-348-TM7c-JB

The next genome examined in this dataset was Candidatus Nanosynbacter HMT-348 strain HMT-348-TM7c-JB (contig_847 in the SC33 metaFlye assembly [Table S3]), which was circular and 841,704 bp in length (Baker, 2022). This Flye draft genome appeared to be the cognate long-read draft genome to the original short-read based draft genome, which was Candidatus_Nanosynbacter_sp._isolate_JCVI_32_bin.19, a 793,808 bp assembly fragmented into 7 contigs (Baker et al., 2021). Following the publication of JB001, JB002, and JB003, the Polypolish short-read polishing tool was published (Wick & Holt, 2022). Polypolish used a novel approach to circumvent the issues in the rRNA and other repeat/conserved regions reported in the paragraph above (Wick & Holt, 2022). Unlike its predecessors, such as Pilon, which map each short read to the best matching location in the draft assembly, Polypolish maps each short read to all matching places in the draft genome (Wick & Holt, 2022). This greatly improves error correction in repeat and conserved regions (Figure 2C) (Wick & Holt, 2022).

To compare the number of errors present in the methods used in the manual pipeline used for JB001, JB002, and JB003 versus Polypolish (i.e. compare the results of the pipelines in Figure 2A and 2C), and determine the optimal pipeline moving forward, each stage of the HMT-348-TM7c-JB assembly was examined for nucleotide identity to the HOMD reference 16S rRNA sequence for HMT-348 (to detect spurious assembly and/or polishing), and was examined for missing or truncated open reading frames (orfs)(mainly due to ONT basecalling errors, which can be corrected by polishing). The Medaka, Flye, and Trycycler assemblies only had one mismatch with the HOMD reference 16S region, while Polypolish had 3, Pilon had 23, Unicycler had 24, and the original Illumina-based SPAdes assembly had 41 (Table 2). This was logical, as the metaSPAdes assembly, using short-reads only, would be expected to have difficulty assembling the 16S region from a metagenomic pool of short reads, and the Pilon and Unicycler (Unicycler itself also uses SPAdes and Pilon as part of its pipeline) polish errors into the rRNA regions due to the conserved regions causing spurious mapping of reads, disrupting variable regions (Figure 2B). It should be noted that the HOMD reference 16S rRNA sequence for HMT-348 is not itself linked to a good quality genome, and there is little knowledge about species and strain diversity within HMT-348, therefore one cannot be sure that the 16S rRNA sequence of HMT-348-TM7c-JB would be expected match the HOMD reference exactly (i.e. the Polypolish, Medaka, Flye, or Trycycler sequences may, in fact, be correct). However, obviously metaSPAdes, Pilon, and Unicycler perform poorly in assembling a correct 16S rRNA gene from a metagenomic read library.

**Table 2:**
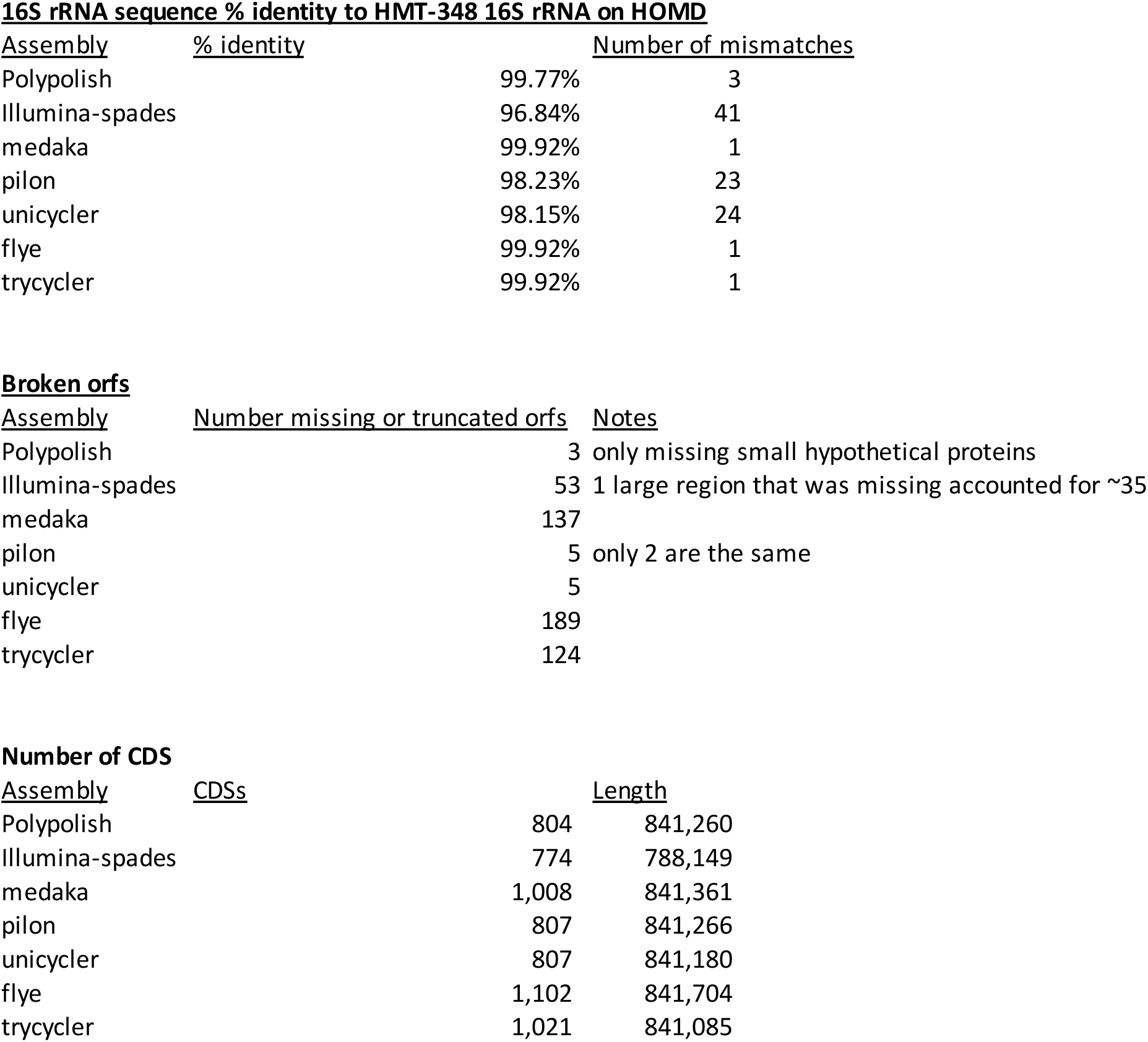
Identifying and comparing errors in HMT-348-TM7c-JB assemblies.

Missing and disrupted open reading frames (orfs) were counted by visualizing the annotated alignment of all 7 assemblies and identifying premature stop codons, split genes, and missing genes. Polypolish performed the best, with only 3 truncated/missing orfs, which were all encoding very small missing hypothetical proteins (which therefore may not even be bona fide genes and errors) (Table 2). The Pilon and Unicycler assemblies each had 5 disrupted orfs (Table 2). The Illumina/metaSPAdes assembly had 53 truncated/missing orfs, 35 of which were missing due to the assembly lacking that entire region (Table 2). As to be expected due to a lack of short-read polishing, the assemblies leaning the most heavily on ONT assembly and polishing did poorly in this regard, with the Medaka, Flye, and Pilon assemblies having 137, 189, and 124 disrupted/missing orfs, respectively (Table 2). The number of disrupted orfs is also apparent when examining the total number of predicted CDSs in each assembly following annotation with Prokka, with the Flye assembly having 298 more ofs than the Polypolish assembly (Table 2). To further validate the metaFlye followed by Polypolish pipeline, JB001 and JB003 were reassembled using metaFlye and Polypolish and compared to the JB001 and JB003 genomes from GenBank which had used the manual Pilon-based approach shown in Figure 2A. JB001 and JB003 quite likely represent different isolates of the same species, and only differed in a few SNPs and two regions that appear to be prophages or other types of mobile elements, which may legitimately be present in only a subset of the metagenomic populations sampled (Baker, 2021b). When compared to the manually polished JB001 and JB003, the Polypolish-polished JB001 and JB003 had >99.99% average nucleotide identity (ANI) of aligned fractions, which had at worst 96.96% alignment percent (AP) of the genome. Polypolish produced identical sequences to the manually polished genomes in one of the 16S rRNA genes and had only one mismatch in JB001 and 3 in JB003 in the second 16S rRNA gene, indicating once again that it largely alleviates the issues observed with the older Pilon polishing tool, which had dozens of mismatches to the HOMD reference. Outside of the 16S mismatches, there were 13 other differences in the assemblies; two were the large putative mobile elements, which were noted previously (Baker, 2021b), and the other 11 were limited to homopolymeric tracts and short repeats, as might be expected from nanopore assembly. Overall, assembly with metaFlye, followed by short-read polishing using Polypolish combined the “best of both worlds” in terms of accuracy, and was used to polish the remaining complete draft genomes described below. The manual polishing approach used on JB001, JB002, and JB003 was also not considered as a viable option moving forward, as it is not amenable to high throughput and more subject to human error. Since HMT-348-TM7c-JB and all other genomes reported here were assembled using the same methods, the following sections will discuss details of the novel genomes themselves.

### *Saccharibacteria* G1-HMT-348 strain HMT-348-TM7c-JB

As recently described (Baker, 2022), HMT-348-TM7c-JB was 841,302 bp. This genome represents the first complete genome from *Saccharibacteria* clade G1 HMT-348, which is one of the most common members of *Saccharibacteria* present in supra-and subgingival plaque, and on the buccal mucosa (McLean et al., 2020; Shaiber et al., 2020). The first draft genome from this clade, TM7c, was published in 2007 (Marcy et al., 2007), however that assembly was later found to contain a significant amount of contamination. More recent studies have binned draft genomes of HMT-348 out of oral metagenomes or single-cell amplified genomes (SAGs), however they were still fragmented into >10 contigs. The exception was the original Illumina-only assembly of HMT-348-TM7c-JB, which was fragmented into 7 contigs (Baker et al., 2021). All of these draft genomes were still significantly incomplete or contaminated (Baker et al., 2021; Cross et al., 2019; Shaiber et al., 2020). Note that the species-level genome bin (SGB) of HMT-348-TM7c-JB in the original metagenomics study also contained two other genomes, likely representing the same species with an ANI of >95% compared to HMT-348-TM7c-JB (Baker et al., 2021). To better examine phylogeny of HMT-348-TM7c-JB and HMT-348, a pangenome including HMT-348-TM7c-JB and 39 other complete *Saccharibacteria* genomes (all complete non-duplicate *Saccharibacteria* genomes on NCBI as of April 2022) was created using Anvi’o (Eren et al., 2021). Only 12 single-copy core genes were common to all 40 genomes. To minimize the effect of gaps on phylogeny, the minimum geometric homogeneity index was set to 0.95, and a maximum functional homogeneity index was set to 0.85 to ensure nearly identical protein sequences were not used (as described in the pangenomics tutorial at <u>anvio.org</u>). This left 4 genes, the ribosomal protein subunits L6 and L27, SecG, and a peptide deformylase, with which to perform a bespoke phylogenetic comparison. Concatenated protein sequences of these 4 genes were used to construct a phylogenetic tree of all 40 complete Saccharibacteria genomes on NCBI, which illustrated that HMT-348-TM7c-JB has its own fairly distinct branch (Figure 3A). To examine unique features of HMT-348-TM7c-JB compared to other clades of oral Saccharibacteria, a more focused pangenome was constructed comparing HMT-348-TM7c-JB with 7 other complete genome representatives from HMT352, HMT-952, HMT-957, HMT-488, HMT-955, and HMT-349 (Figure 3B). The analysis identified 218 pan-G1 Core Genes, 327 genes which were unique to HMT-348-TM7c-JB, and 21 genes which were missing in HMT-348-TM7c-JB, but present in all other clades (Figure 3B, pangenome summary table available at: https://github.com/jonbakerlab/nanopore-oral-genomes). Note that the pangenome data tables generated by this study are too large (>50MB, in some cases) to be included as supplemental material, and therefore have been made accessible on GitHub. Interestingly, many of the features missing in HMT-348-TM7c-JB were either chaperones or involved in DNA repair, including *xseA, uvrD* and *dnaK*. In addition to the well conserved Type IV and Type II secretion systems encoded by all the G1 Saccharibacteria genomes, HMT-348-TM7c-JB appeared to have its own distinct Type VI and II secretion systems. Further examination of the HMT-348 genome and comparison to other Saccharibacteria will provide insight into the host-epiboint dynamics and ecological role of this abundant yet enigmatic and uncultivated bacterial taxa.

**Figure 3:**
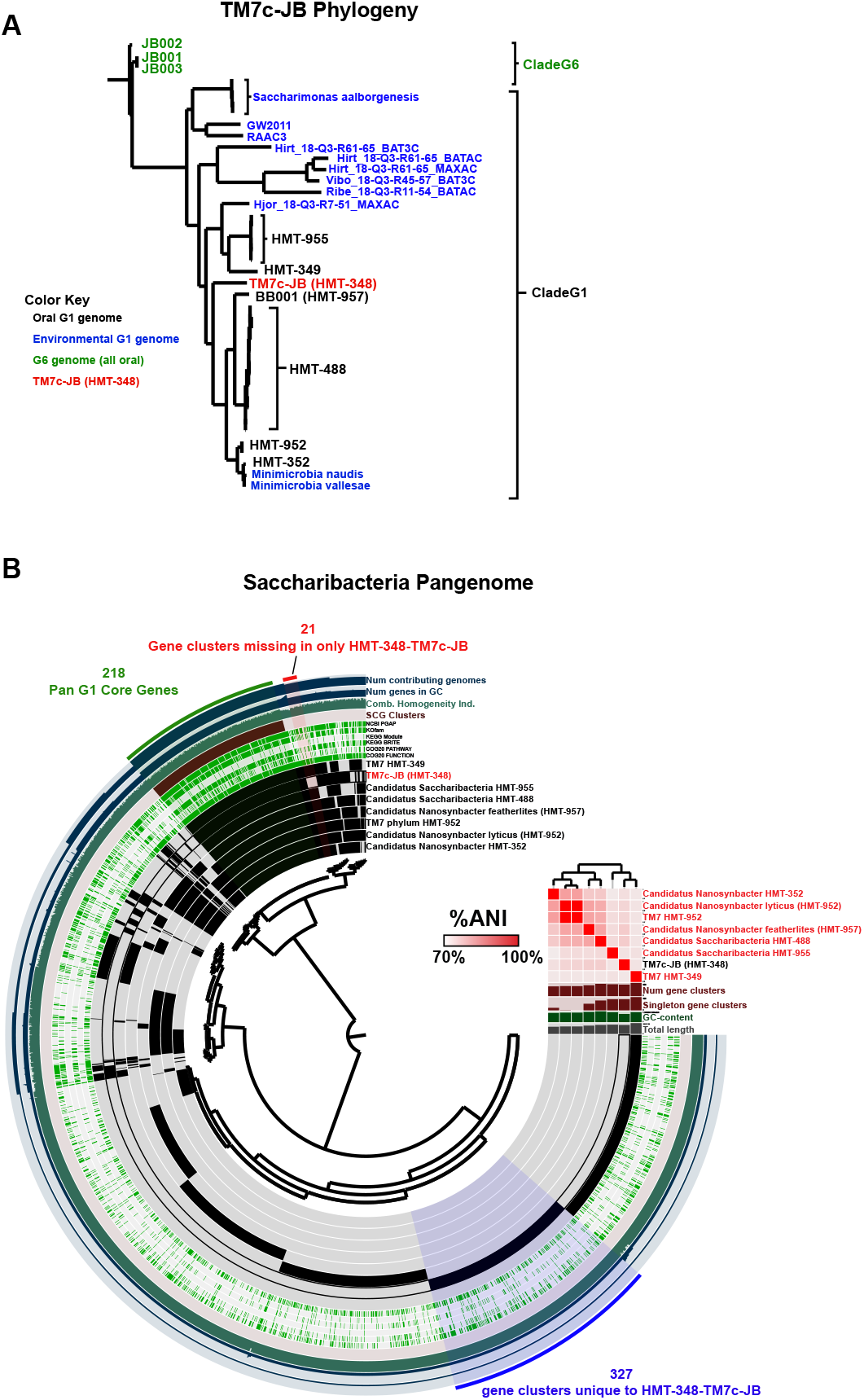
Phylogeny and pangenome of *Saccharibacteria* and HMT-348-TM7c-JB. **(A)** Updated phylogeny of complete *Saccharibacteria* genomes. Phylogenetic tree based upon concatenated protein sequences of 4 single-copy core genes with optimal geometric and functional homogeneity indices. All complete *Saccharibacteria* genomes on GenBank (April 2022) were included, and the clade G6 genomes were used to root the tree (all remaining genomes were clade G1). To improve readability, several species-level clades were collapsed to one label. Clade G6 genomes, which are all from human oral sources, are labeled in green, human oral G1 genomes are labeled in black (except HMT-348-TM7c-JB), environmental G1 genomes are labeled in blue, and HMT-348-TM7c-JB is labeled in red. **(B)** Clade G1 *Saccharibacteria* pangenome. The dendrogram in the center organizes the 2,545 gene clusters identified across the genomes represented by the innermost 8 layers: TM7 HMT-349, HMT-348-TM7c-JB (HMT-348), Candidatus *Saccharibacteria* HMT-955, Candidatus *Saccharibacteria* HMT-488, Candidatus *Nanosynbacter featherlites* (HMT957), TM7 HMT-952, Candidatus *Nanosynbacter lyticus* (HMT-952), and *Candidatus Nanosynbacter* HMT-352. The data points within these 8 layers indicate the presence of a gene cluster in a given genome. From inside to outside, the next 6 layers indicate known vs unknown COG function, COG pathway, KEGG Brite, KEGG module, KOfam, NCBI PGAP annotation. The next 4 layers indicate single copy core gene (SCG) clusters, the combined homogeneity index, the number of genes in the gene cluster, and the number of contributing genomes. The outmost layer indicates the gene clusters present in the following groups: Genes missing in only HMT-348, Pan-G1 Core Genes, and genes unique to HMT-348. The 8 genome layers are ordered based on the tree of the %ANI comparison, which is displayed with the red and white heat map. The layers underneath the %ANI heat map, from top to bottom, indicate: number of gene clusters, number of singleton gene clusters, GC content, and total length of each genome.

#### Actinomyces graevenitzii

There were two genomes of *Actinomyces graevenitzii* assembled by this study, JCVI-JB-Ag32 from SC33 at 2,088,213 bp and JCVI-JB-Ag28 from SC28 at 2,122,055 bp. These two genomes had an ANI of 97.03% across an AP of 93.97%. JCVI-JB-Ag28 had very poor coverage of Illumina reads (0.2X), while JCVI-JB-Ag32 had an Illumina coverage of 68.5X, which was excellent. Therefore, JCVI-JB-Ag28 genome likely contained many nanopore errors, undoubtedly contributing to the ANI difference between the genomes. JCVI-JB-Ag32 had a 16S rRNA gene with 99.963% identity (4 mismatches) to *Actinomyces graevenitzii* HMT-866 strain F0530 (from HOMD), while JCVI-JB-Ag28 had 99.344% identity (5 mismatches) to the same reference 16S rRNA. There are currently no complete genomes of *Actinomyces graevenitzii* available on HOMD or NCBI, with the two draft genomes on HOMD being fragmented into 10 and 29 contigs with a 2.09-2.21 Mbp size. The two genomes from this study had an ANI of 95-96% against the HOMD genomes with an AP of 91-92%. In the previous metagenomics study, there were 2 other *A. graevenitzii* in the SGB ANI >95% SGB with the draft genomes of JCVI-JB-Ag28 and JCVI-JB-Ag32 (Baker et al., 2021). Pangenome analysis of JCVI-JB-Ag32 and 26 *Actinomyces* complete genomes from GenBank identified 655 gene clusters unique to *A. graevenitzii* and 8 gene clusters missing in only *A. graevenitzii* (Figure 4, pangenome summary table available at: https://github.com/jonbakerlab/nanopore-oral-genomes). JCVI-JB-Ag28 was not included in the pangenome analysis due to the likely high rate of ONT errors, which would affect gene calling. The pangenome analysis indicated that *A. naeslundii* GCA002355915 is likely misidentified, as it was much more closely related to *A. oris*, rather than other genomes of *A. naeslundii*, in terms of both %ANI and gene cluster presence/absence (Figure 4). The pangenome analysis also indicated that *A. graevenitzii* was comparatively more divergent from other *Actinomyces*, with %ANI < 76% compared to all other genomes examined. *A. graevenitzii* is frequently isolated from the oral cavity (Wang et al., 2019), occasionally from the gut (Ruigrok et al., 2021), and also causes infections of the lung (Gliga et al., 2014). Recently, *A. graevenitzii* was shown to inhibit the growth *Streptococcus* and *Staphylococcus* while also co-operating with *Staphylococcus aureus* to evade neutrophil attack (Jalali et al., 2021). Deeper examination of the genes present in *A. graevenitzii* will give insight into its metabolic capabilities and ecological role.

**Figure 4:**
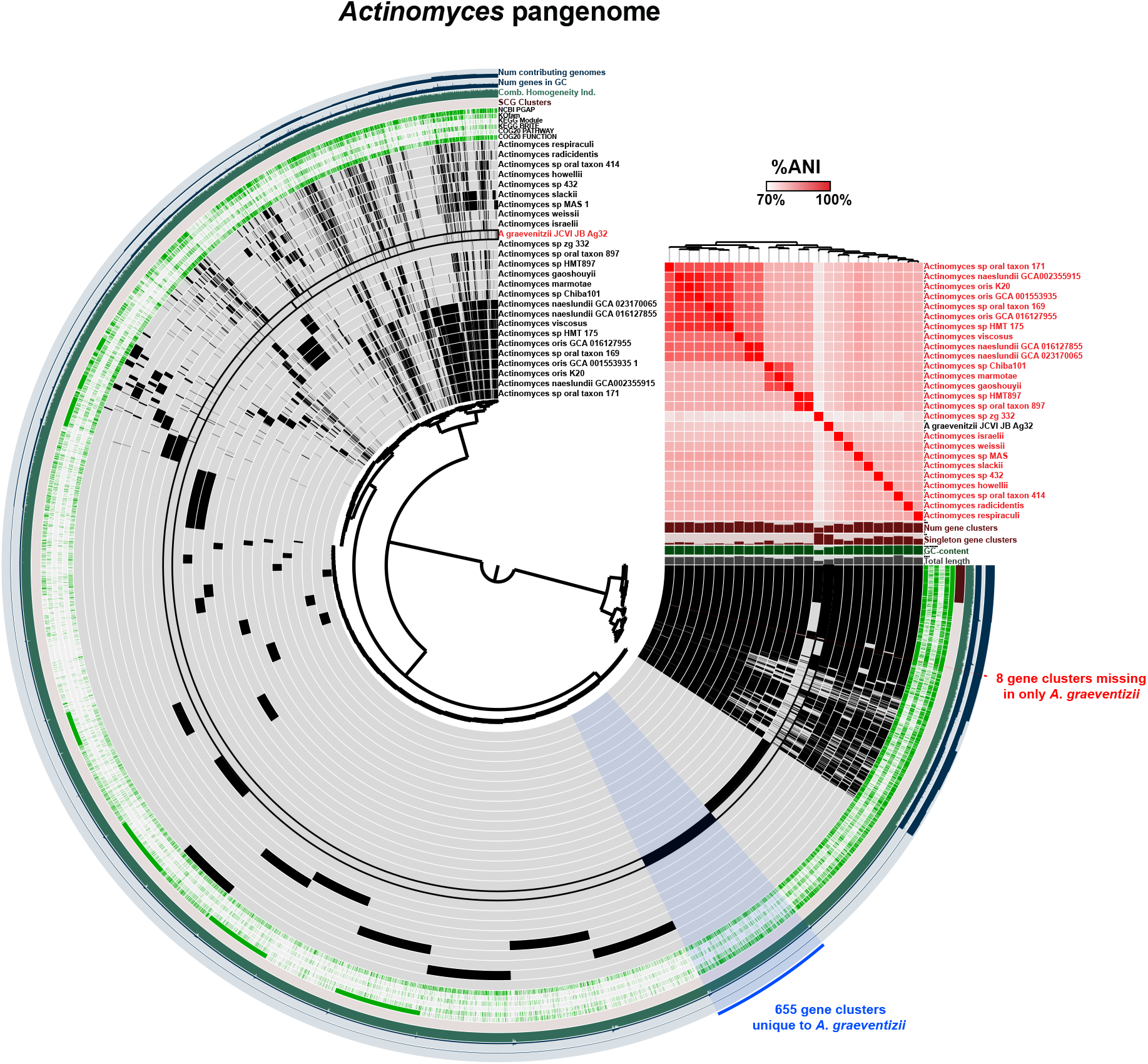
The *Actinomyces* pangenome. The dendrogram in the center organizes the 12,168 gene clusters identified across the indicated genomes represented by the innermost 26 layers. The data points within these 26 layers indicate the presence of a gene cluster in a given genome. From inside to outside, the next 6 layers indicate known vs unknown COG function, COG pathway, KEGG Brite, KEGG module, KOfam, and NCBI PGAP annotation. The next 4 layers indicate single copy core gene (SCG) clusters, the combined homogeneity index, the number of genes in the gene cluster, and the number of contributing genomes. The outmost layer indicates the gene clusters present in the following groups: Gene clusters missing in only *A. graevenitzii*, and gene clusters unique to *A. graevenitzii*. The 26 genome layers are ordered based on the tree of the %ANI comparison, which is displayed with the red and white heat map. The layers underneath the %ANI heat map, from top to bottom, indicate: number of gene clusters, number of singleton gene clusters, GC content, and total length of each genome.

#### Rothia mucilaginosa

There were two complete genomes of *Rothia mucilaginosa* obtained in this study, JCVI-JB-Rm27 from SC27 at 2,258,635 bp and 16.6X Illumina coverage, and JCVI-JB-Rm28 from SC28 at 2,259,930 bp with 5.3X Illumina coverage. Compared to each other, the two genomes obtained in this study have an ANI of 95.22% and an AP of 95.58%. Both strains have a 16S rRNA gene with 99.775% identity (3 mismatches) to *Rothia mucilaginosa* HMT-681 reference strain, DY-18. There are currently 4 complete genomes of *Rothia mucilaginosa* on 4 on NCBI. Compared to these other *R. mucilaginosa* genomes, 27 and 28 have ANIs of ∼93% and 90-92% AP. This is lower than might be expected, and may indicate the presence of multiple subspecies or genospecies within *R. mucilaginosa*. In the previous metagenomic study, there were 13 other *Rothia* bins in the SGB with the JCVI-JB-Rm27 and JCVI-JB-Rm28 draft genomes, with 11 having of these having an ANI of >95% to JCVI-JB-Rm27 and JCVI-JB-Rm28 (Baker et al., 2021). Pangenome analysis was performed using JCVI-JB-Rm27 and 17 other complete *Rothia* genomes from GenBank (Figure 5, pangenome summary table available at: https://github.com/jonbakerlab/nanopore-oral-genomes). This analysis identified 1,291 gene clusters unique to *R. mucilaginosa* and 27 gene clusters missing only in *R. mucilaginosa* (Figure 5). Compared to the other complete *R. mucilaginosa* genomes, JCVI-JB-Rm27 had 51 unique gene clusters and there were 27 gene clusters missing in only JCVI-JB-Rm27 (Figure 5). Although *Rothia* are known to cause various types of opportunistic infections (Fatahi-Bafghi, 2021), they have also received attention recently as a potential probiotic in the context of dental caries (Rosier, Moya-Gonzalvez, Corell-Escuin, & Mira, 2020), as they and other nitrate-reducing oral bacteria have been associated with good dental health (Agnello et al., 2017; Baker et al., 2021).

**Figure 5:**
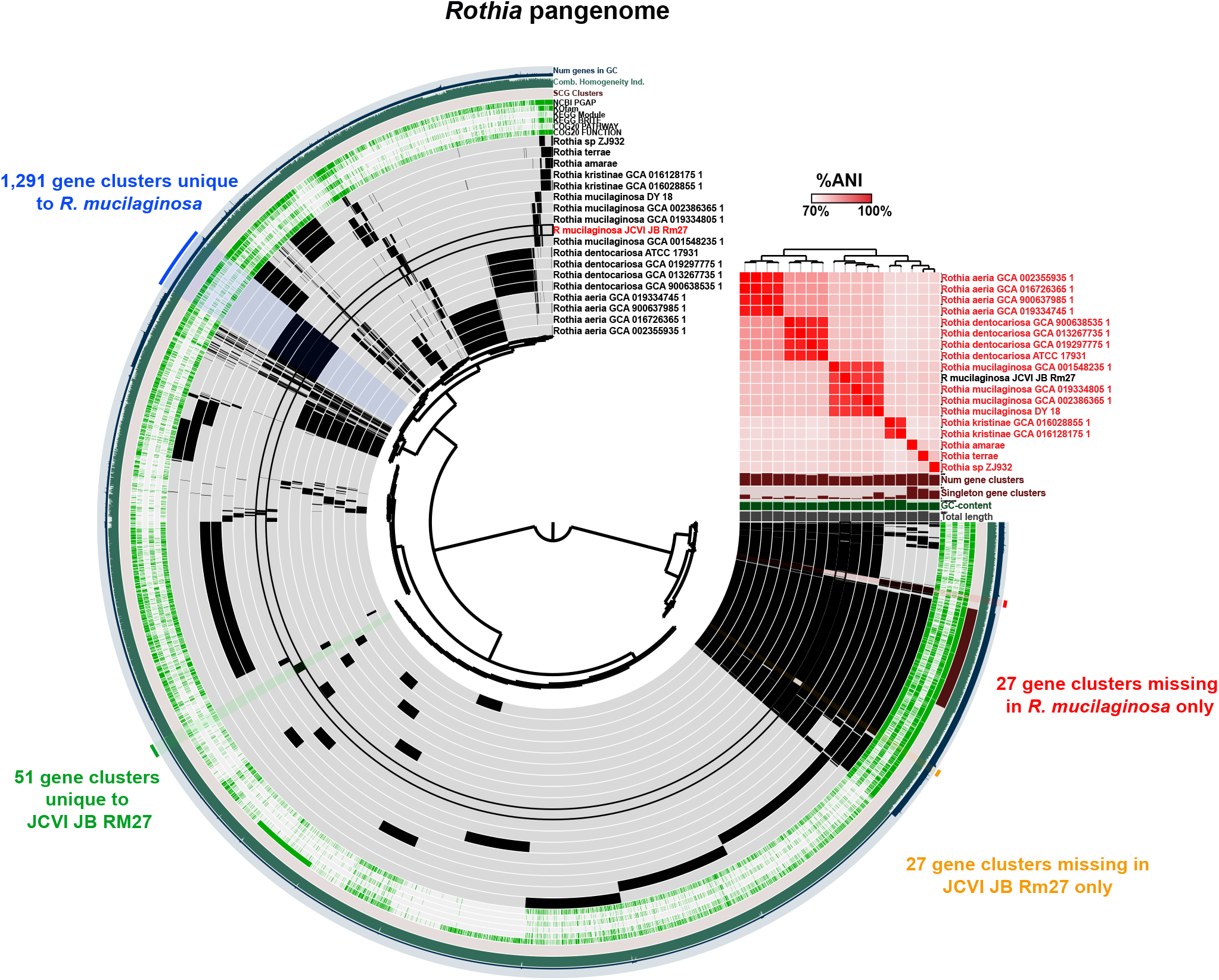
The *Rothia* pangenome. The dendrogram in the center organizes the 8.118 gene clusters identified across the indicated genomes represented by the innermost 18 layers. The data points within these 18 layers indicate the presence of a gene cluster in a given genome. From inside to outside, the next 6 layers indicate known vs unknown COG function, COG pathway, KEGG Brite, KEGG module, KOfam, and NCBI PGAP annotation. The next 4 layers indicate single copy core gene (SCG) clusters, the combined homogeneity index, the number of genes in the gene cluster, and the number of contributing genomes. The outmost layer indicates the gene clusters present in the following groups: Gene clusters missing in only *R. mucilaginosa*, gene clusters unique to *R. mucilaginosa*, gene clusters unique to JCVI-JB-Rm27, and gene clusters missing in only JCVI-JB-Rm27. The 18 genome layers are ordered based on the tree of the %ANI comparison, which is displayed with the red and white heat map. The layers underneath the %ANI heat map, from top to bottom, indicate: number of gene clusters, number of singleton gene clusters, GC content, and total length of each genome.

#### Mogibacterium diversum

From SC33, a circular 1,770,007 bp genome of *Mogibacterium diversum*, JCVI-JB-Md32, was obtained. The 16S rRNA gene from JCVI-JB-Md32 had 100% identity to *Mogibacterium diversum* HMT-593 strain ATCC 700923. JCVI-JB-Md32 had an ANI of 96.08% and AP of 89.40% to *M. diversum* strain CCUG47132, the only other complete *M. diversum* genome on HOMD and NCBI. This genome had excellent coverage with the Illumina reads at 837X. *M. diversum* belongs to the very poorly understood family, Eubacteriales Family XIII, *Insertae Sedis*, which also contains the prevalent oral resident, [Eubacterim] *sulci*. Interestingly, the JCVI-JB-Md32 draft genome had ANI >95% with 10 other genomes in the same SGB in the previous study (Baker et al., 2021). Having 10 genomes of this species independently assembled and binned from a study of 47 human subjects indicates that this taxon is rather prevalent, despite being relatively unstudied. Pangenome analysis of JCVI-JB-Md32 and all 6 other complete genomes within this family was performed (Figure 6, pangenome summary table available at: https://github.com/jonbakerlab/nanopore-oral-genomes). This analysis identified 743 gene clusters unique to *M. diversum* and 162 gene clusters missing in only *M. diversum*. Compared to CCUG47132, JCVI-JB-Md32 had 163 unique gene clusters and only 3 gene clusters missing (Figure 6). This complete genome and pangenome data will be a useful resource in the study of *M. diversum*, as only 1 study of this common member of the oral flora has been reported (Nakazawa et al., 2002).

**Figure 6:**
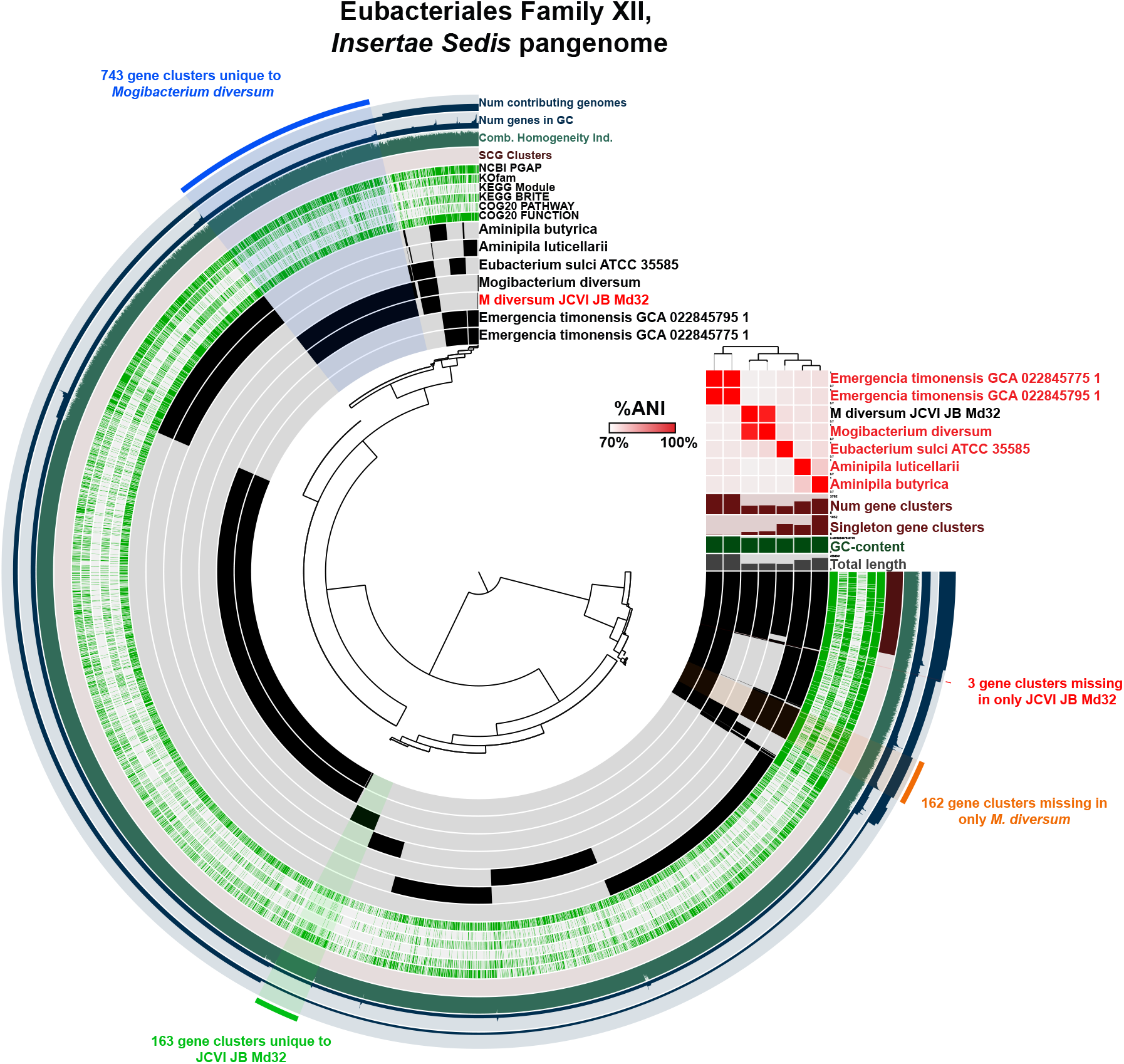
The Eubacteriales Family XIII, Insertae Sedis pangenome. The dendrogram in the center organizes the 8.084 gene clusters identified across the indicated genomes represented by the innermost 7 layers. The data points within these 7 layers indicate the presence of a gene cluster in a given genome. From inside to outside, the next 6 layers indicate known vs unknown COG function, COG pathway, KEGG Brite, KEGG module, KOfam, and NCBI PGAP annotation. The next 4 layers indicate single copy core gene (SCG) clusters, the combined homogeneity index, the number of genes in the gene cluster, and the number of contributing genomes. The outmost layer indicates the gene clusters present in the following groups: Gene clusters missing in only *M. diversum*, gene clusters unique to *M. diversum*, gene clusters unique to JCVI-JB-Md32, and gene clusters missing in only JCVI-JB-Md32. The 7 genome layers are ordered based on the tree of the %ANI comparison, which is displayed with the red and white heat map. The layers underneath the %ANI heat map, from top to bottom, indicate: number of gene clusters, number of singleton gene clusters, GC content, and total length of each genome.

#### Lancefieldella parvula

JCVI-JB-Lp32, a complete genome of *Lancefieldella parvula* (formerly known as *Atopobium parvulum*) was 1,624,536 bp. The JCVI-JB-Lp32 16S rRNA gene is 99.544% identical (6 mismatches) to *Lancefieldella parvula* HMT-723 strain ATCC22793. One complete genome of *L. parvula*, IPP1246, is available and was published in 2009 (Copeland et al., 2009). JCVI-JB-Lp32 and IPP1246 have an ANI of 88.7% over an AP of 84.13%. It is interesting that although the genome assembled here and the NCBI reference genome have an extremely high 16S rRNA identity, and certainly are the same species based upon 16S rRNA sequence, that the overall genome ANI is significantly lower than 95%, which is typically expected for genomes from the same species. Notably, although the ANI to IPP1246 genome was lower, in the original metagenomics study with the Illumina draft assembly of JCVI-JB-Lp32, there were 22 genomes in the same SGB with ANI > 95% to JCVI-JB-Lp32 (Baker et al., 2021). This indicates that perhaps these 22 draft genomes and JCVI-JB-Lp32 are a distinct subspecies compared to IP1246, and that *L. parvula* has a relatively plastic genome, or that perhaps these genomes are a separate species entirely. Pangenome analysis of JCVI-JB-Lp32 and 6 other Atopobiaceae genomes identified 327 gene clusters unique to *L. parvula* and 62 gene clusters missing in only *L. parvula* (Figure 7, pangenome summary table available at: https://github.com/jonbakerlab/nanopore-oral-genomes). Compared to IPP1246, JCVI-JB-Lp32 had 178 unique gene clusters, and was missing 50 gene clusters uniquely compared to all other Atopobiaceae analyzed (Figure 7). In the oral cavity, *L. parvulum* has been associated with occlusal lesions in caries (Fakhruddin, Samaranayake, Hamoudi, Ngo, & Egusa, 2022), or with a healthy microbiome, in the context of periodontal disease (Kumar et al., 2003). In the gut, *L. parvulum* has been associated with colorectal cancer (Yachida et al., 2019) and the onset of Crohn’s disease, due to its ability to produce H2S through the SufS cysteine desulfurase (Karunakaran et al., 2022; Mottawea et al., 2016).

**Figure 7:**
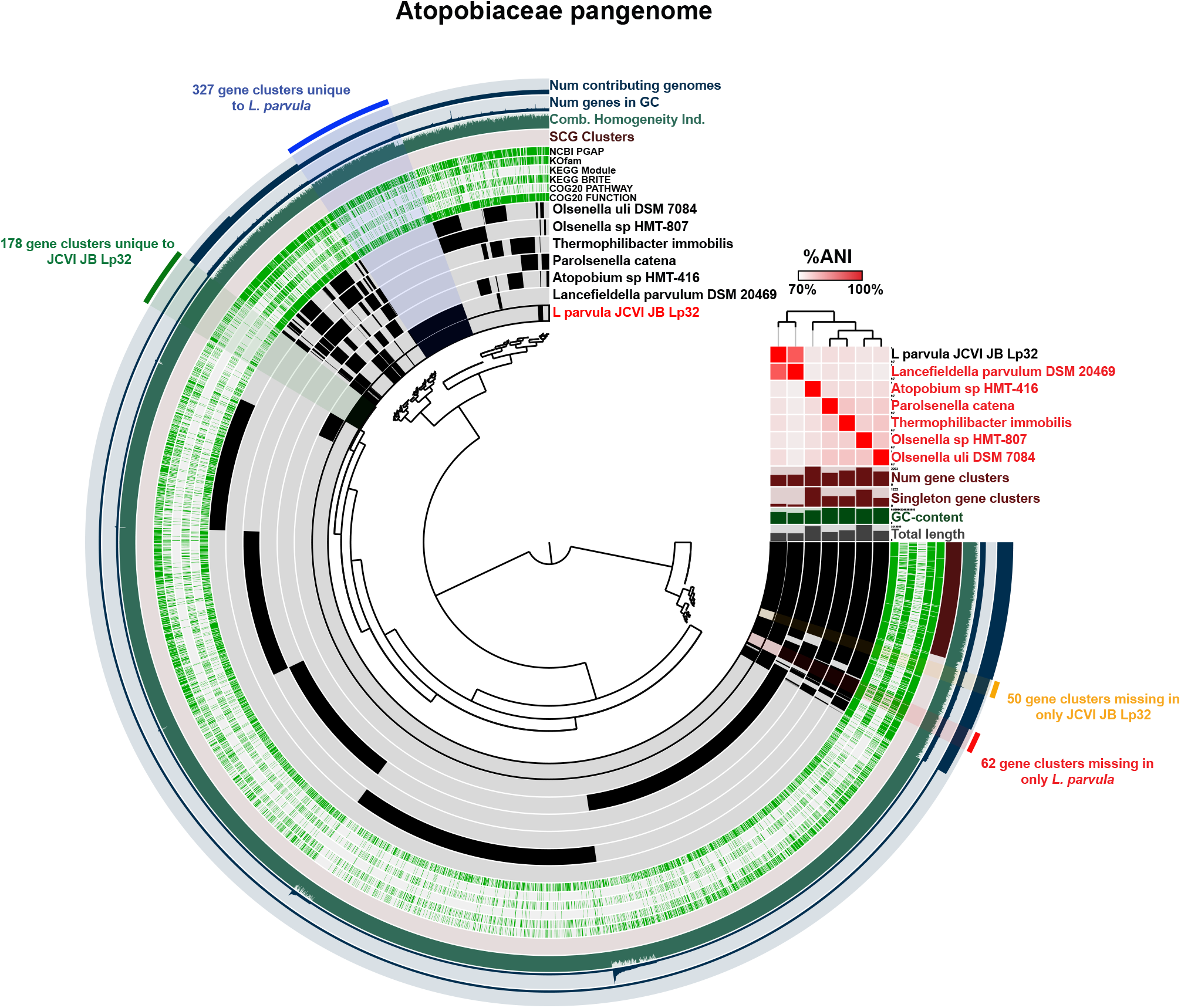
The Atopobiceae pangenome. The dendrogram in the center organizes the 6,488 gene clusters identified across the indicated genomes represented by the innermost 7 layers. The data points within these 7 layers indicate the presence of a gene cluster in a given genome. From inside to outside, the next 6 layers indicate known vs unknown COG function, COG pathway, KEGG Brite, KEGG module, KOfam, and NCBI PGAP annotation. The next 4 layers indicate single copy core gene (SCG) clusters, the combined homogeneity index, the number of genes in the gene cluster, and the number of contributing genomes. The outmost layer indicates the gene clusters present in the following groups: Gene clusters missing in only *L. parvula*, gene clusters unique to *L. parvula*, gene clusters unique to JCVI-JB-Lp32, and gene clusters missing in only JCVI-JB-Lp32. The 7 genome layers are ordered based on the tree of the %ANI comparison, which is displayed with the red and white heat map. The layers underneath the %ANI heat map, from top to bottom, indicate: number of gene clusters, number of singleton gene clusters, GC content, and total length of each genome.

### Lachnospiraeceae HMT-096

JCVI-JB-L28 was assembled from SC28, and was 2,401,457 bp, and had a 16S rRNA gene seq is 99.7% identical (3 mismatches) to Lachnospiraceae [G-2] HMT-096 from HOMD. There is one complete genome from Lachnospiraceae [G-2] HMT-096 on NCBI, CP073340, deposited in 2021 by The Forsyth Institute, which has an ANI of 98.585% to the JCVI-JB-L28 over an AP of 87.79%. As this genome did not have good coverage by the Illumina reads (0.3X coverage), this assembly currently represents a draft genome that was not further analyzed, and likely would have a higher ANI to the published genome with increased short-read coverage for polishing. Lachnospiraceae HMT-096 is very poorly understood, but its abundance in saliva was positively correlated with hemodialysis in patients with chronic and end-stage kidney disease (Duan et al., 2020).

### Ruminococcaceae HMT-075 chr1

The 1,207,213 bp JCVI-JB-R28 Ruminococcaceae chromosome obtained had a 16S rRNA gene with 99.773 % identity (3 mistmatches) to Ruminococcaceae [G-1] bacterium HMT-075 Clone F058. There are currently no genomes of this taxa on HOMD or NCBI. Due to the small size of this genome compared to other Ruminococcaceae, which typically have 2-4 Mbp genomes, and the fact that some other Ruminococcaceae genomes have two chromosomes (Wegmann et al., 2014), it is likely that this represents a single chromosome of a taxon with multiple chromosomes. Phylophlan3 (Asnicar et al., 2020) was used to examine all other large contigs (>700,000 bp, linear and circular), from SC28, to determine if they may represent the other chromosomes of this taxon, however none were predicted to be from the same taxon. Although this is only an incomplete genome due to the fact that there is only one chromosome, it does represent the first genome sequence to be associated with the 16S sequence for Ruminococcaceae HMT-075, a taxon which almost nothing is known about other than its existence.

### Conclusions

This study serves as proof-of-concept, and provides a protocol, for obtaining complete genomes from metagenomes derived from human saliva using complementary ONT and Illumina sequencing. This represents a major advance, as obtaining complete genomes previously required isolation of the microbe and/or manual PCR/sequencing steps due to repeat regions. In this manner, obtaining complete genomes, which include the 16S rRNA regions that are typically not accurately assembled and binned from Illumina reads alone, will finally provide genomes for the large number of taxa that are currently only associated with 16S rRNA sequences. Although the greatly reduced error rate of current ONT technologies, and the process of polishing using Illumina reads, results in highly accurate genomes, it is likely they still have higher error rates than genomes produced with Illumina sequencing where the gaps/repeats are manually sequenced using Sanger technologies. However, the greatly increased throughput of the methods presented here, and lack of need for laboratory isolation and growth of the organisms, more than offset the disadvantage in sequence accuracy. Deeper sequencing and improved methods for obtaining ultra-high molecular weight gDNA are likely to produce even more circular assemblies per sample. The information provided by the genomes obtained here will be useful to help determine the metabolic capabilities, ecological roles, and pathogenic potential of the cognate bacterial species, particularly those that have not been isolated or cultivated.

## Materials and methods

### DNA extraction

As described in the Results section, several methods were attempted to optimize extraction of HMW gDNA. [1] the Chen and Burne phenol:chloroform method described in (Y. Y. Chen et al., 1996) [2] the open-source Bio-On-Magnetic-Beads (BOMB) gDNA Extraction using guanidine isothiocyanate (GITC) lysis and purification using silane magnetic beads (detailed protocol available at bomb.bio), [3] the lysis steps from the Chen and Burne protocol (Y. Y. Chen et al., 1996), followed by purification using silane beads (instead of phenol:chloroform), and [4] the Monarch Genomic DNA Purification Kit from New England Biolabs (using manufacturer’s instructions). Approach [1] gave the best combination of yield and size of HMW gDNA, and is therefore reported in full detail here (gDNA chromatograms and concentrations from all 4 methods is available in Supplemental Files 1 and 2): 545 μl saliva (from a frozen aliquot) was added to 545 μl TE buffer (100 mM, 20 mM) containing 10 mg/ml lysozyme and 300 U/ml mutanolysin, and was incubated for 30 min at 37°C. 100 μl 20% SDS was added, and the sample was incubated at 65°C for 15 min. 300 μl TE (100 mM, 20 mM) was added, and the whole volume of lysate was transferred to a 2 ml screwcap tube containing 0.1 mm glass beads. The sample was homogenized for 30 s on a FastPrep-24 homogenizer (MP Biomedicals) and the lysate (none of the foam) was transferred to a new 1.5 microcentrifuge tube and cooled to 37° C. 2 μl of proteinase K was added and the sample was incubated for 30 min at 37° C. 100 μl of 5M NaCl was added, followed by 80 μl of 10 CTAB in 0.7M NaCl which had been warmed to 65° C. The sample was then incubated at 65° C for 20 min. 750 μl phenol:chloroform:isoamyl alcohol (25:24:1) was added, the solution was mixed by inversion and then centrifuged at 12,000 x g for 1 minute. The aqueous phase was extracted and extractions were repeated until the white interface between the aqueous and organic layers was gone (typically 2-3 exactions). The aqueous phase was then extracted once with 750 μl chloroform:isoamyl alcohol (24:1). 750 μl ice-cold 100% isopropanol was added and the sample was centrifuged at 12,000 x g for 30 min. The pellet was washed with 70% ethanol then rinsed with 100% ethanol. The pellet was then resuspended in 100 μl water. An additional final ethanol precipitation step was performed. 10 μl 3M sodium acetate pH 5.2 and 250 μl of ice-cold 100% ethanol were added to the sample and the sample was centrifuged at 12,000 x g for 10 minutes at 4°C. The supernatant was decanted, then 250 μl of 70% was added, and the sample was centrifuged for 5 minutes at 12,000 x g. The supernatant was decanted and the pellet was dried in a speed-vac. The final pellet was resuspended in 50 μl of molecular grade water. The quality of the DNA was checked using a TapeStation (Agilent Technologies, Santa Clara, CA, USA), and Qubit Fluorimeter (Thermo Fisher Scientific, Waltham, MA, USA).

### Illumina sequencing

The short-read sequencing was performed as previously described (Baker et al., 2021). Briefly, libraries were generated using the Nextera XT DNA Library Preparation Kit (Illumina, Inc., San Diego, CA, USA) using the manufacturer’s instructions and the sequencing run was performed on a NextSeq500 (Illumina, Inc., San Diego, CA, USA).

### Nanopore sequencing

ONT sequencing was performed as previously described (Baker, 2021a, 2021b; Baker & Edlund, 2020). Briefly, the long-read library was prepared using a Ligation Sequencing Kit (Oxford Nanopore Technologies, Oxford, UK) and sequenced on a GridION using an R9.4.1 flow cell (Oxford Nanopore Technologies, Oxford, UK). Base calling, quality control, and adapter trimming were performed using Guppy v4.0.11/MinKNOW v20.06.9 (Oxford Nanopore Technologies, Oxford, UK).

### Genome assembly. (JB001, JB002, JB003)

Two independent methods generated improved draft assemblies (compared to the draft assemblies published in Baker et al. 2021 (Baker et al., 2021). (i) Human reads were removed from the long-read assemblies using minimap2 v2.17-r941 (Li, 2018), and the remaining long reads were assembled using metaFlye v2.8-b1674 (Kolmogorov et al., 2020). MegaBLAST v2.2.26 (Zhang, Schwartz, Wagner, & Miller, 2000) was used to identify the circular contigs of interest within the metagenome assemblies. (ii) Long reads mapping to the draft genomes of JB001, JB002, and JB003 were extracted using minimap2. These long reads, along with the short reads used to generate the original JB001, JB002, and JB003 draft assemblies, were used by Unicycler v0.4.8 (Wick et al., 2017) to obtain draft genomes. Short contigs in the Unicycler assemblies were removed based on disparate GC content, coverage, and BLAST hits to other organisms (Anvi’o v7-dev) (Eren et al., 2021), leaving single circular contigs. Trycycler v0.3.0 (Wick et al., 2021) was used to develop consensus assemblies from the draft assemblies. The resulting assemblies were polished using Medaka v1.0.3 (https://github.com/nanoporetech/medaka), then Pilon v1.23 (Walker et al., 2014). Circulator v1.5.5 (Hunt et al., 2015) was used to rotate the genome sequences such that the start sites were at the *dnaA* gene. Default parameters were used unless otherwise noted. JB001, JB002, and JB003 were annotated initially using Prokka (Seemann, 2014), while the final genomes submitted to NCBI were annotated using the NCBI Prokaryotic Genome Annotation Pipeline v5.1. As noted in above in the Results and in (Baker, 2021b), further examination indicated the Medaka and Pilon polishing steps introduced errors into the rRNA regions, but properly removed errors from other locations in the genomes. Therefore, the rRNA versions were manually corrected to those obtained from the Flye assemblies. **(TM7c-JB)** The same approaches just detailed for JB001, JB002, and JB003 were also used for HMT-348-TM7c-JB. In addition, Polypolish (v0.4.3) (Wick & Holt, 2022), which was published during preparation of this study was attempted in parallel on the HMT-348-TM7c-JB contig from the metaFlye assembly. The Polypolish tool utilized the short reads from the cognate short-read libraries which had mapped to the metaFlye draft contig. 16S rRNA sequences were compared using the HOMD 16S rRNA Sequence Identification tool (https://www.homd.org/refseq/refseq_blastn). Disrupted and missing orfs were identified, and %ANI and %AP were calculated using CLC Genomics Workbench v21.0.3 (Qiagen, Inc., Maryland, USA). The results obtained from metaFlye followed by Polypolish closely resembled those from the final composite methods used in the JB001, JB002, and JB003 final assemblies (i.e. rRNA regions from metaFlye, remainder of the genome from Unicycler/Flye/Trycycler/Medaka/Pilon), but was much less computationally and temporally expensive (Table 2). Therefore, this pipeline, metaFlye, followed by Polypoish was used for all other genomes in this study. **(all other genomes)** Short reads were mapped to the metaFlye draft genomes using BWA-MEM (Li & Durbin, 2009) and used by Polypolish for error correction. The start sequences were rotated to *dnaA* using Circulator and annotated using NCBI PGAP.

### Phylogenomics and pangenomics

The Anvi’o (v-dev) pangenomics workflow (Delmont & Eren, 2018; Eren et al., 2015; Eren et al., 2021) was implemented using Snakemake (Koster & Rahmann, 2012) and used to perform the pangenomics analysis. For the phylogenetic analysis of *Saccharibacteria*, HMT-348-TM7c-JB and HMT-348, a pangenome including HMT-348-TM7c-JB and 39 other complete *Saccharibacteria* genomes (all complete non-duplicate *Saccharibacteria* genomes on NCBI as of April 2022) was created. Only 12 single-copy core genes were common to all 40 genomes. To minimize the effect of gaps on phylogeny, the minimum geometric homogeneity index was set to 0.95, and a maximum functional homogeneity index was set to 0.85 to ensure nearly identical protein sequences were not used. This left 4 genes, the ribosomal protein subunits L6 and L27, SecG, and a peptide deformylase. Concatenated protein sequences of these 4 genes were used to construct a phylogenetic tree of all 40 complete *Saccharibacteria* genomes on NCBI using Anvi’o.

## Acknowledgements

I thank Karrie Goglin-Alemeida, Jelena Jablanovic, and Kara Riggsbee for performing the library preparation and sequencing. This research was supported by NIH/NIDCR K99-DE029228.

## Data Availability

The short reads used to generate the assemblies were published previously (Baker et al., 2021) and are available in the SRA database with the accession numbers SRX4318838, SRX4318837, and SRX4318835. The long reads used to assemble the metagenomes are available in the SRA database with accession numbers SRX15103396, SRX15103397, and SRX15103398. The complete genomes of JCVI-JB-Rm27, JCVI-JB-Ag32, JCVI-JB-Md32, JCVI-JB-Lp32, and HMT-348-TM7c-JB are available on NCBI GenBank as accession numbers CP097094, CP097095, CP097093, CP097092, and CP090820, respectively. Since JCVI-JB-L28, JCVI-JB-Ag28, and JCVI-JB-Rm28 have low Illumina coverage and are likely to contain ONT errors, and JCVI-JB-R28-chr1, is likely not a complete genome, these sequences are available at https://github.com/jonbakerlab/nanopore-oral-genomes. The other draft assemblies of HMT-348-TM7c-JB (metaSPAdes, Unicycler, Trycycler, Medaka, Pilon, and metaFlye) and the Polypolish versions of JB001 and JB003 are also available at https://github.com/jonbakerlab/nanopore-oral-genomes. The pangenome data tables are too large (some >50MB) to be included as Supplemental Material, and therefore have been made available at https://github.com/jonbakerlab/nanopore-oral-genomes.

## Figure Legends

**Table S1: Summary of the contigs produced by metaFlye assembly of SC27**.

**Table S2: Summary of the contigs produced by metaFlye assembly of SC28**.

**Table S3: Summary of the conrigs produced by metaFlye assembly of SC33**.

**File S1: TapeStation report of HMW gDNA concentration and quality using gDNA extraction methods [1-3] as described in Materials and Methods**. Method [1] the Chen and Burne phenol:chloroform method described in (Y. Y. Chen et al., 1996): -bead-beating = well D1 / +bead-beating = well E1. Method [2] the open-source Bio-On-Magnetic-Beads (BOMB) gDNA Extraction using guanidine isothiocyanate (GITC) lysis and purification using silane magnetic beads (detailed protocol available at bomb.bio): -bead-beating = well F1 / + bead-beating = well G1. Method [3] the lysis steps from the Chen and Burne protocol (Y. Y. Chen et al., 1996), followed by purification using silane beads (instead of phenol:chloroform): -bead-beating = well B1 / + bead-beating = well C1.

**File S2: TapeStation report of HMW gDNA concentration and quality using gDNA extraction method [4] as described in Materials and Methods**. Method [4] the Monarch Genomic DNA Purification Kit from New England Biolabs (using manufacturer’s instructions): -bead-beating = well B1 / + bead-beating = well C1.

