## Supplementary material for "Using nanopore sequencing to obtain complete genomes from saliva samples": File S2

### Filename: 2020-08-20-01_JIRA_1517_gDNA.gDNA

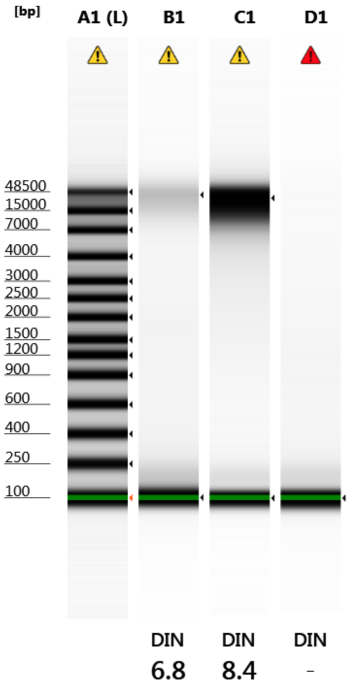


Default image (Contrast 100%)

#### Sample Info

| Well | DIN | Conc. [ng/µl] | Sample Description | Alert | Observations |
|---|---|---|---|---|---|
| A1 | - | 61.8 | Ladder | ⚠ | Caution! Expired ScreenTape device; Ladder |
| B1 | 6.8 | 4.21 | JIRA 1517 S. mutans gDNA 7 | ⚠ | Caution! Expired ScreenTape device; Sample concentration outside functional range for DIN |
| C1 | 8.4 | 20.0 | JIRA 1517 S. mutans gDNA 8 | ⚠ | Caution! Expired ScreenTape device |
| D1 | - | 2.02 | H2O | ⚠ | Caution! Expired ScreenTape device; Sample concentration outside functional range for DIN |


© Agilent Technologies, Inc. 2017

#### A1: Ladder

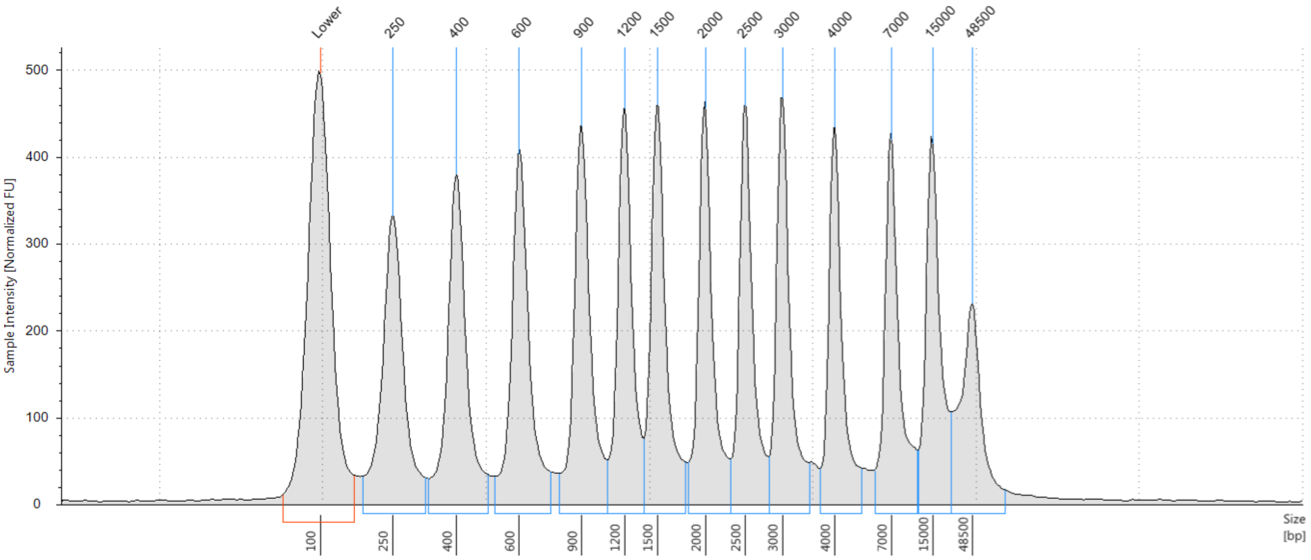


##### Sample Table

| Well | DIN | Conc. [ng/µl] | Sample Description | Alert | Observations |
|---|---|---|---|---|---|
| A1 | - | 61.8 | Ladder | ⚠ | Caution! Expired ScreenTape device; Ladder |

##### Peak Table

| Size [bp] | Calibrated Conc. [ng/µl] | Assigned Conc. [ng/µl] | % Integrated Area | From [bp] | To [bp] | Peak Comment |
|---|---|---|---|---|---|---|
| 100 | 8.50 | 8.50 | - | 63 | 154 | |
| 250 | 5.09 | - | 8.35 | 171 | 318 | |
| 400 | 5.11 | - | 8.39 | 325 | 490 | |
| 600 | 5.12 | - | 8.41 | 513 | 735 | |
| 900 | 5.12 | - | 8.40 | 777 | 1070 | |
| 1200 | 4.87 | - | 8.00 | 1070 | 1375 | |
| 1500 | 4.98 | - | 8.17 | 1375 | 1777 | |
| 2000 | 4.79 | - | 7.86 | 1807 | 2308 | |
| 2500 | 4.57 | - | 7.50 | 2308 | 2817 | |
| 3000 | 4.62 | - | 7.57 | 2817 | 3478 | |
| 4000 | 4.24 | - | 6.96 | 3701 | 5218 | |
| 7000 | 4.38 | - | 7.19 | 5918 | 11234 | |
| 15000 | 4.42 | - | 7.26 | 11533 | 21110 | |
| 48500 | 3.46 | - | 5.68 | 21110 | >60000 | |
| - | - | - | - | - | - | |

 © Agilent Technologies, Inc. 2017 

#### B1: JIRA 1517 S. mutans gDNA 7

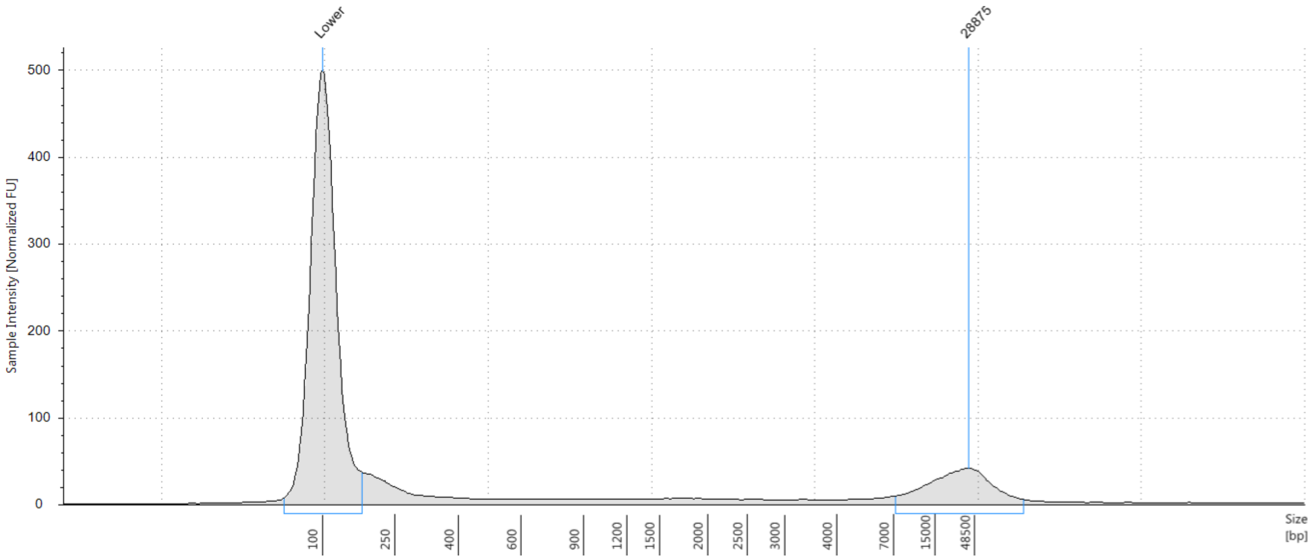


##### Sample Table

| Well | DIN | Conc. [ng/µl] | Sample Description | Alert | Observations |
|---|---|---|---|---|---|
| B1 | 6.8 | 4.21 | JIRA 1517 S. mutans gDNA 7 | ⚠ | Caution! Expired ScreenTape device; Sample concentration outside functional range for DIN |

##### Peak Table

| Size [bp] | Calibrated Conc. [ng/µl] | Assigned Conc. [ng/µl] | % Integrated Area | From [bp] | To [bp] | Peak Comment |
|---|---|---|---|---|---|---|
| 100 | 8.50 | 8.50 | - | 61 | 163 | |
| 28875 | 1.68 | - | 50.20 | 7059 | >60000 | |
| >60000 | 0.137 | - | 4.09 | >60000 | >60000 | |
| - | - | - | - | - | - | |

 © Agilent Technologies, Inc. 2017 

#### C1: JIRA 1517 S. mutans gDNA 8

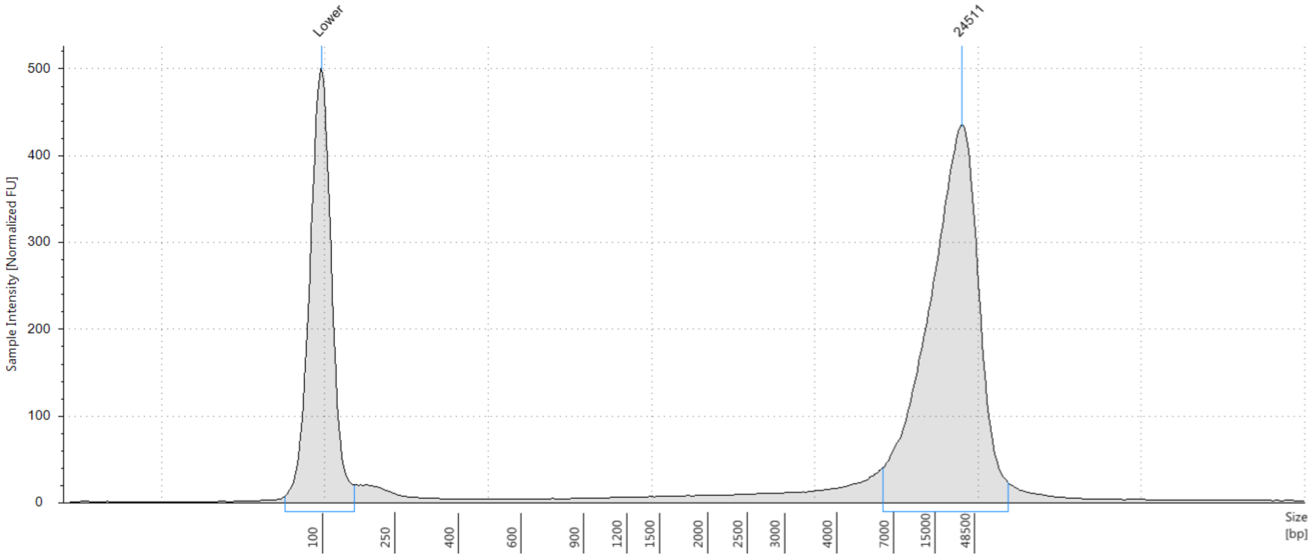


##### Sample Table

| Well | DIN | Conc. [ng/µl] | Sample Description | Alert | Observations |
|---|---|---|---|---|---|
| C1 | 8.4 | 20.0 | JIRA 1517 S. mutans gDNA 8 | ⚠ | Caution! Expired ScreenTape device |

##### Peak Table

| Size [bp] | Calibrated Conc. [ng/µl] | Assigned Conc. [ng/µl] | % Integrated Area | From [bp] | To [bp] | Peak Comment |
|---|---|---|---|---|---|---|
| 100 | 8.50 | 8.50 | - | 63 | 151 | |
| 24511 | 16.8 | - | 93.51 | 6267 | >60000 | |
| - | - | - | - | - | - | |

 © Agilent Technologies, Inc. 2017 

## D1: H2O

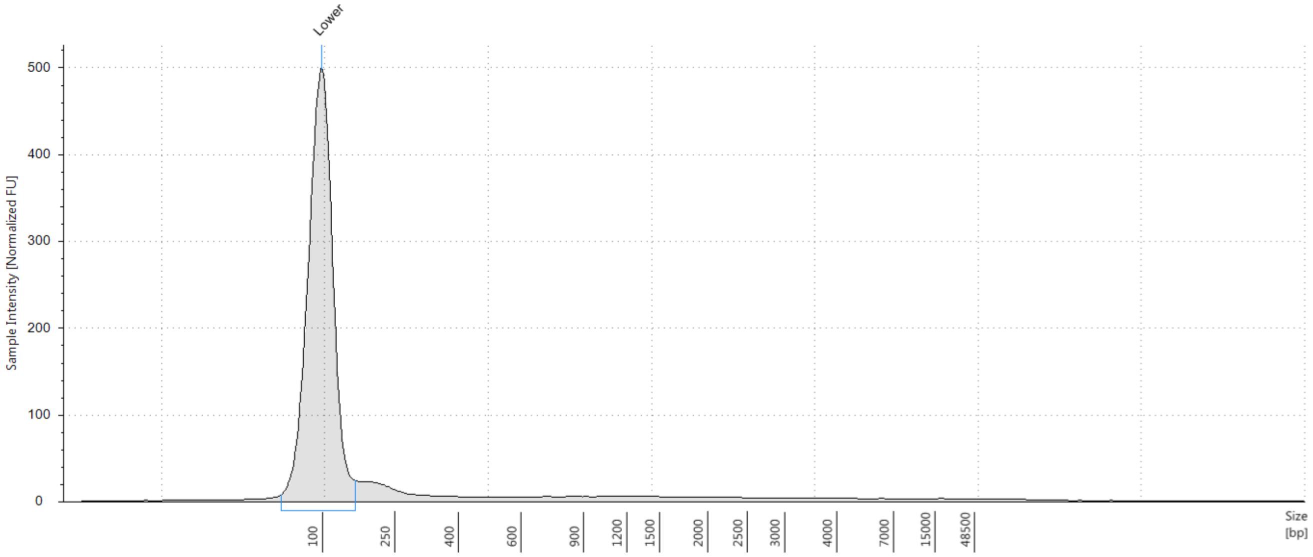


##### Sample Table

| Well | DIN | Conc. [ng/µl] | Sample Description | Alert | Observations |
|---|---|---|---|---|---|
| D1 | - | 2.02 | H2O | ⚠ | Caution! Expired ScreenTape device; Sample concentration outside functional range for DIN |

##### Peak Table

| Size [bp] | Calibrated Conc. [ng/µl] | Assigned Conc. [ng/µl] | % Integrated Area | From [bp] | To [bp] | Peak Comment |
|---|---|---|---|---|---|---|
| 100 | 8.50 | 8.50 | - | 60 | 153 | |


© Agilent Technologies, Inc. 2017
