## Supplementary material for "Using nanopore sequencing to obtain complete genomes from saliva samples": File S1

### Filename: 2020-08-17-01_JIRA_1518.gDNA

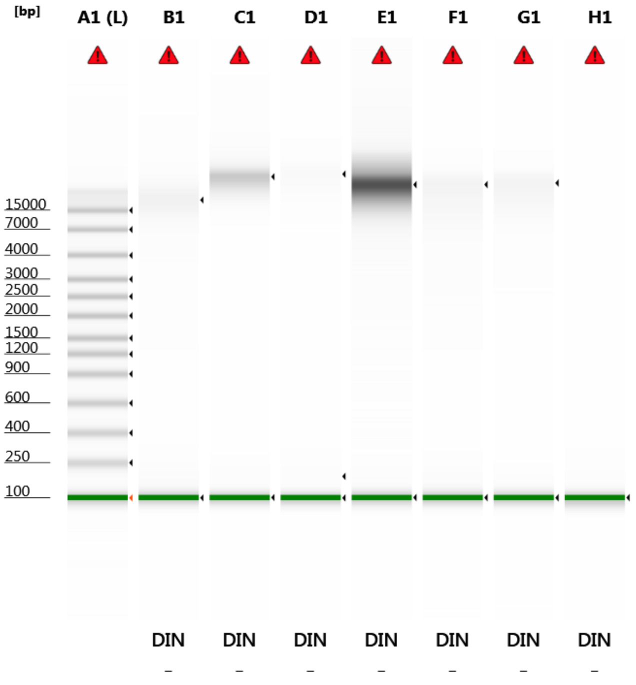


Default image (Contrast 100%)

#### Sample Info

| Well | DIN | Conc. [ng/µl] | Sample Description | Alert | Observations |
|---|---|---|---|---|---|
| A1 | - | 67.4 | Ladder | ⚠ | Issue with ladder peak detection (too few peaks detected); Caution! Expired ScreenTape device; Ladder |
| B1 | - | 10.1 | JIRA 1518 1 | ⚠ | Issue with ladder peak detection (too few peaks detected); Caution! Expired ScreenTape device |
| C1 | - | 18.7 | JIRA 1518 2 | ⚠ | Issue with ladder peak detection (too few peaks detected); Caution! Expired ScreenTape device |
| D1 | - | 6.78 | JIRA 1518 3 | ⚠ | Issue with ladder peak detection (too few peaks detected); Caution! Expired ScreenTape device; Sample concentration outside recommended range |
| E1 | - | 90.2 | JIRA 1518 4 | ⚠ | Issue with ladder peak detection (too few peaks detected); Caution! Expired ScreenTape device |
| F1 | - | 11.0 | JIRA 1518 5 | ⚠ | Issue with ladder peak detection (too few peaks detected); Caution! Expired ScreenTape device |
| G1 | - | 8.40 | JIRA 1518 6 | ⚠ | Issue with ladder peak detection (too few peaks detected); Caution! Expired ScreenTape device; Sample concentration outside recommended range |
| H1 | - | 1.46 | EB buffer | ⚠ | Issue with ladder peak detection (too few peaks detected); Caution! Expired ScreenTape device; Sample concentration outside functional range for DIN |

 © Agilent Technologies, Inc. 2017 

#### A1: Ladder

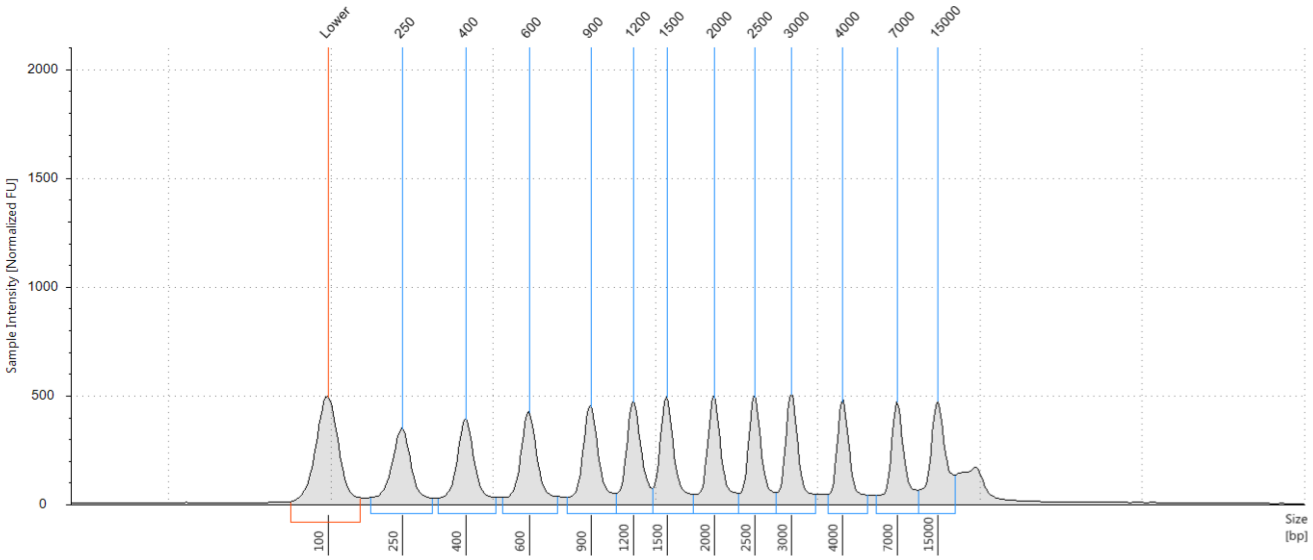


**Sample Table**

| Well | DIN | Conc. [ng/µl] | Sample Description | Alert | Observations |
|---|---|---|---|---|---|
| A1 | - | 67.4 | Ladder | ⚠ | Issue with ladder peak detection (too few peaks detected); Caution! Expired ScreenTape device; Ladder |

**Peak Table**

| Size [bp] | Calibrated Conc. [ng/µl] | Assigned Conc. [ng/µl] | % Integrated Area | From [bp] | To [bp] | Peak Comment |
|---|---|---|---|---|---|---|
| 100 | 8.50 | 8.50 | - | 24 | 165 | |
| 250 | 5.26 | - | 8.49 | 185 | 320 | |
| 400 | 5.28 | - | 8.51 | 333 | 493 | |
| 600 | 5.39 | - | 8.70 | 516 | 736 | |
| 900 | 5.40 | - | 8.72 | 784 | 1080 | |
| 1200 | 5.33 | - | 8.60 | 1080 | 1375 | |
| 1500 | 5.32 | - | 8.58 | 1375 | 1779 | |
| 2000 | 5.27 | - | 8.51 | 1779 | 2293 | |
| 2500 | 5.02 | - | 8.11 | 2293 | 2788 | |
| 3000 | 4.95 | - | 7.99 | 2788 | 3472 | |
| 4000 | 4.62 | - | 7.46 | 3722 | 5385 | |
| 7000 | 4.71 | - | 7.60 | 5846 | 11138 | |
| 15000 | 5.31 | - | 8.57 | 11138 | 18310 | |
| - | - | - | - | - | - | |

 © Agilent Technologies, Inc. 2017 

### B1: JIRA 1518 1

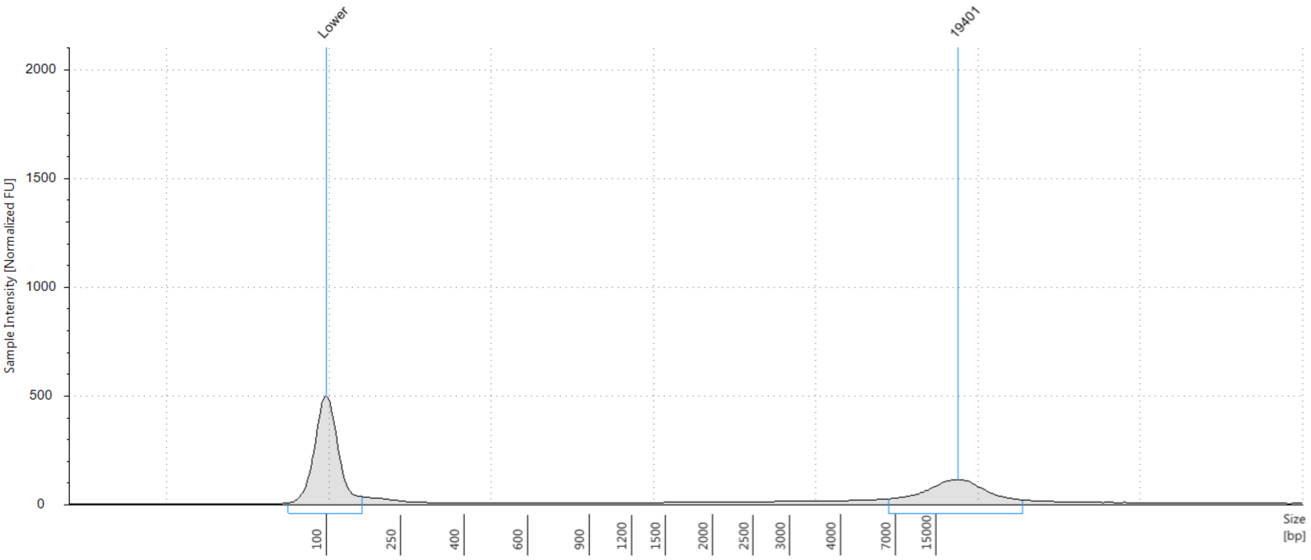


**Sample Table**

| Well | DIN | Conc. [ng/µl] | Sample Description | Alert | Observations |
|---|---|---|---|---|---|
| B1 | - | 10.1 | JIRA 1518 1 | ⚠ | Issue with ladder peak detection (too few peaks detected); Caution! Expired ScreenTape device |

**Peak Table**

| Size [bp] | Calibrated Conc. [ng/µl] | Assigned Conc. [ng/µl] | % Integrated Area | From [bp] | To [bp] | Peak Comment |
|---|---|---|---|---|---|---|
| 100 | 8.50 | 8.50 | - | 23 | 171 | |
| 19401 | 5.18 | - | 96.75 | 6664 | 32175 | |
| - | - | - | - | - | - | |

 © Agilent Technologies, Inc. 2017 

#### C1: JIRA 1518 2

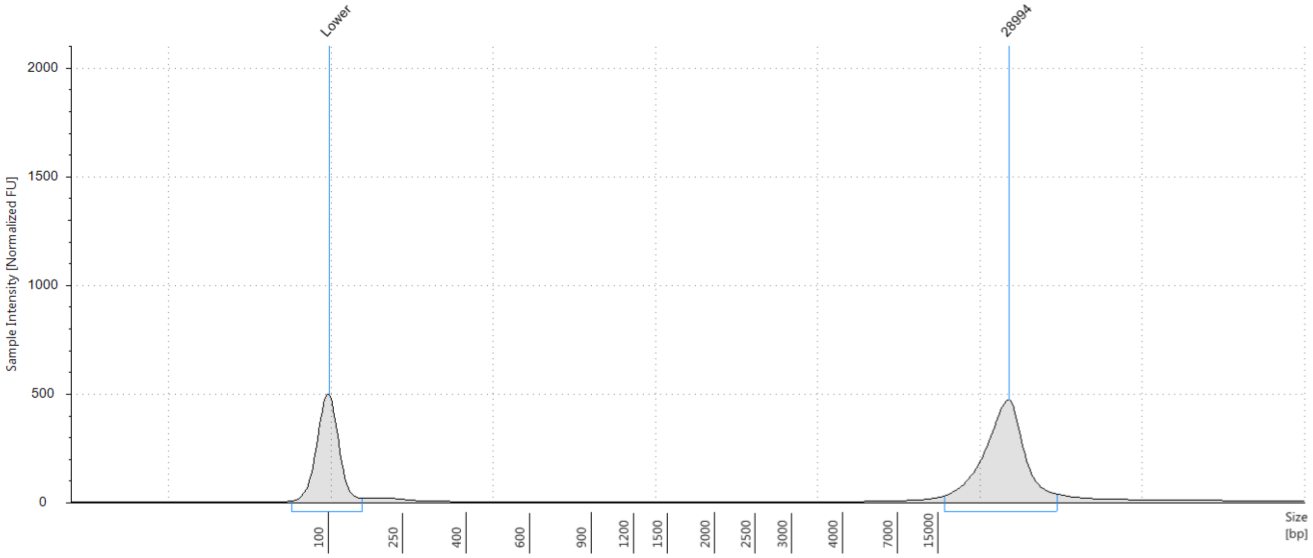


**Sample Table**

| Well | DIN | Conc. [ng/µl] | Sample Description | Alert | Observations |
|---|---|---|---|---|---|
| C1 | - | 18.7 | JIRA 1518 2 | ⚠ | Issue with ladder peak detection (too few peaks detected); Caution! Expired ScreenTape device |

**Peak Table**

| Size [bp] | Calibrated Conc. [ng/µl] | Assigned Conc. [ng/µl] | % Integrated Area | From [bp] | To [bp] | Peak Comment |
|---|---|---|---|---|---|---|
| 100 | 8.50 | 8.50 | - | 24 | 167 | |
| 28994 | 14.4 | - | 98.08 | 16245 | 38343 | |
| - | - | - | - | - | - | |


© Agilent Technologies, Inc. 2017

#### D1: JIRA 1518 3

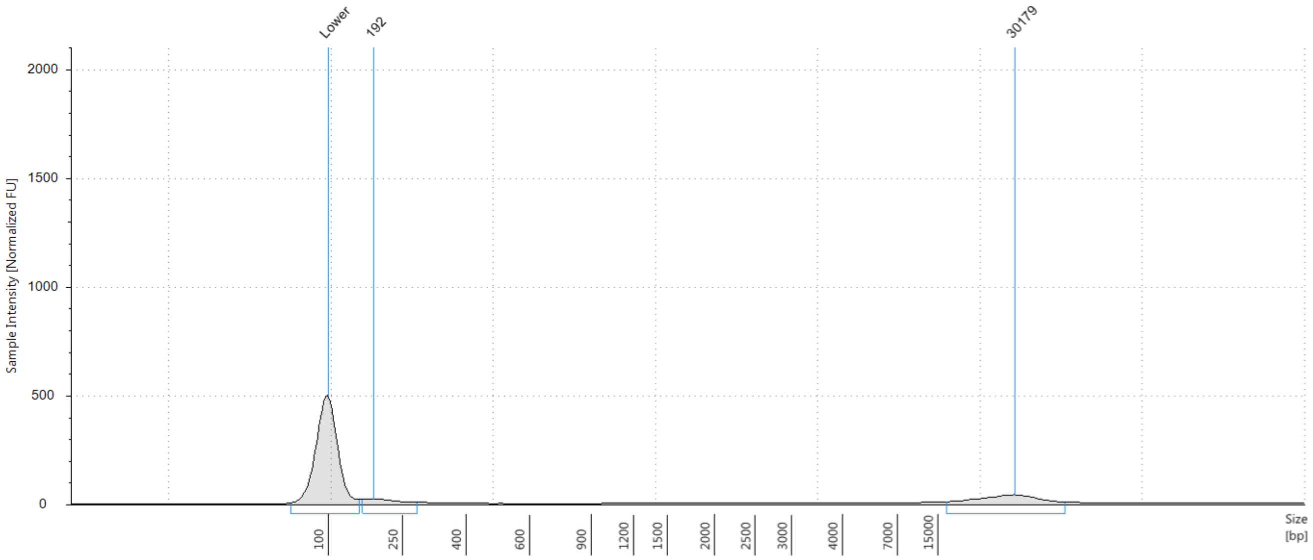


##### Sample Table

| Well | DIN | Conc. [ng/µl] | Sample Description | Alert | Observations |
|---|---|---|---|---|---|
| D1 | - | 6.78 | JIRA 1518 3 | ⚠ | Issue with ladder peak detection (too few peaks detected); Caution! Expired ScreenTape device; Sample concentration outside recommended range |

##### Peak Table

| Size [bp] | Calibrated Conc. [ng/µl] | Assigned Conc. [ng/µl] | % Integrated Area | From [bp] | To [bp] | Peak Comment |
|---|---|---|---|---|---|---|
| 100 | 8.50 | 8.50 | - | 25 | 163 | |
| 192 | 0.610 | - | 4.85 | 169 | 283 | |
| 30179 | 2.06 | - | 16.36 | 16717 | 39996 | |
| >60000 | 1.14 | - | 9.07 | >60000 | >60000 | |
| - | - | - | - | - | - | |


© Agilent Technologies, Inc. 2017

### E1: JIRA 1518 4

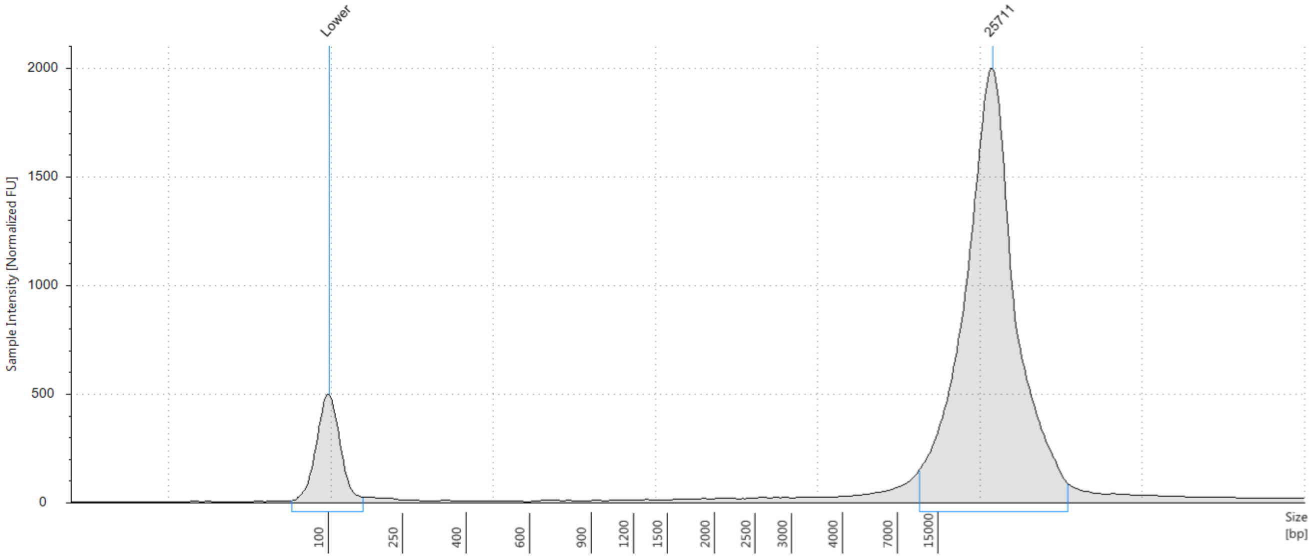


**Sample Table**

| Well | DIN | Conc. [ng/µl] | Sample Description | Alert | Observations |
|---|---|---|---|---|---|
| E1 | - | 90.2 | JIRA 1518 4 | ⚠ | Issue with ladder peak detection (too few peaks detected); Caution! Expired ScreenTape device |

**Peak Table**

| Size [bp] | Calibrated Conc. [ng/µl] | Assigned Conc. [ng/µl] | % Integrated Area | From [bp] | To [bp] | Peak Comment |
|---|---|---|---|---|---|---|
| 100 | 8.50 | 8.50 | - | 25 | 169 | |
| 25711 | 74.1 | - | 51.95 | 11344 | 40643 | |
| >60000 | 6.40 | - | 4.49 | >60000 | >60000 | |
| - | - | - | - | - | - | |

 © Agilent Technologies, Inc. 2017 

#### F1: JIRA 1518 5

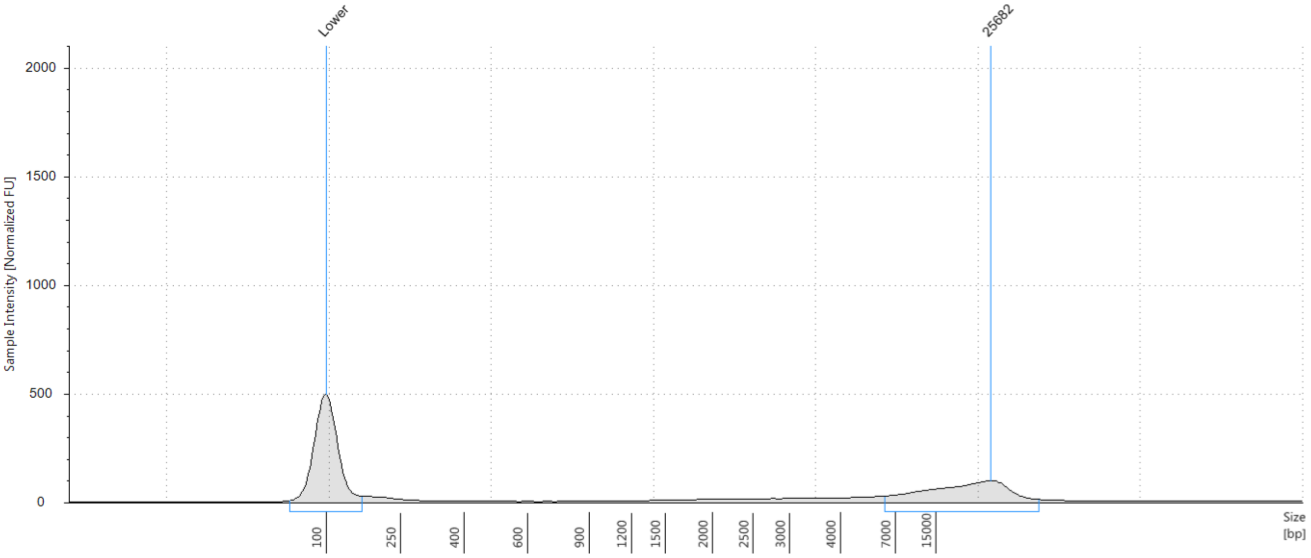


##### Sample Table

| Well | DIN | Conc. [ng/µl] | Sample Description | Alert | Observations |
|---|---|---|---|---|---|
| F1 | - | 11.0 | JIRA 1518 5 | ⚠ | Issue with ladder peak detection (too few peaks detected); Caution! Expired ScreenTape device |

##### Peak Table

| Size [bp] | Calibrated Conc. [ng/µl] | Assigned Conc. [ng/µl] | % Integrated Area | From [bp] | To [bp] | Peak Comment |
|---|---|---|---|---|---|---|
| 100 | 8.50 | 8.50 | - | 26 | 171 | |
| 25682 | 5.52 | - | 83.47 | 6404 | 35224 | |
| - | - | - | - | - | - | |

 © Agilent Technologies, Inc. 2017 

#### G1: JIRA 1518 6

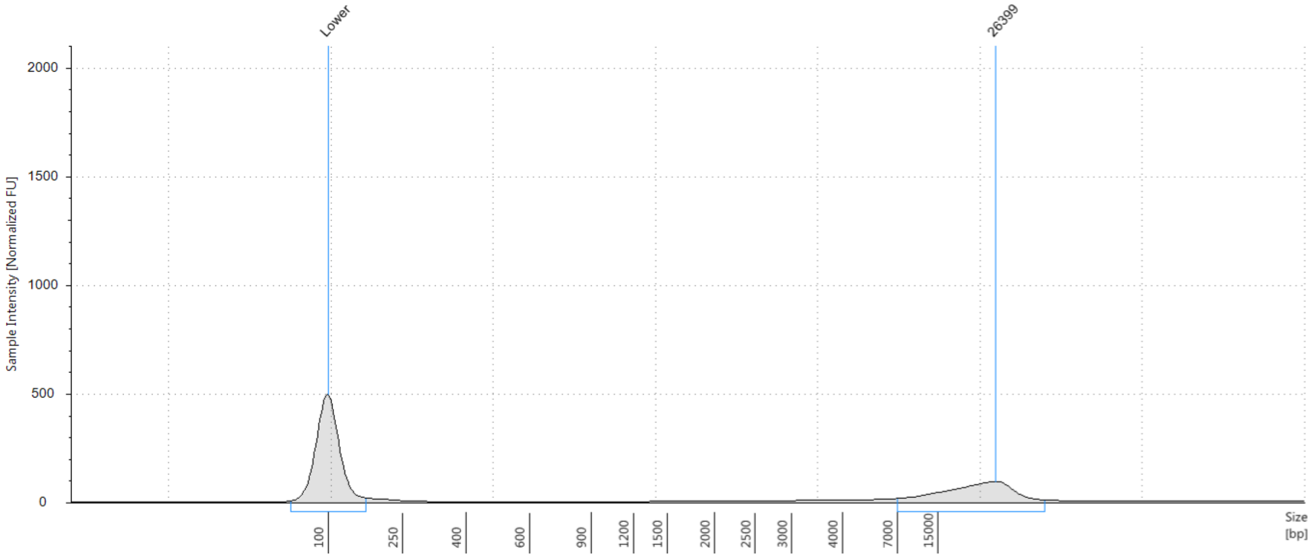


**Sample Table**

| Well | DIN | Conc. [ng/µl] | Sample Description | Alert | Observations |
|---|---|---|---|---|---|
| Gl | - | 8.40 | JIRA 1518 6 | ⚠ | Issue with ladder peak detection (too few peaks detected); Caution! Expired ScreenTape device; Sample concentration outside recommended range |

**Peak Table**

| Size [bp] | Calibrated Conc. [ng/µl] | Assigned Conc. [ng/µl] | % Integrated Area | From [bp] | To [bp] | Peak Comment |
|---|---|---|---|---|---|---|
| 100 | 8.50 | 8.50 | - | 24 | 176 | |
| 26399 | 4.69 | - | 74.61 | 7073 | 35914 | |
| - | - | - | - | - | - | |


© Agilent Technologies, Inc. 2017

### H1: EB buffer

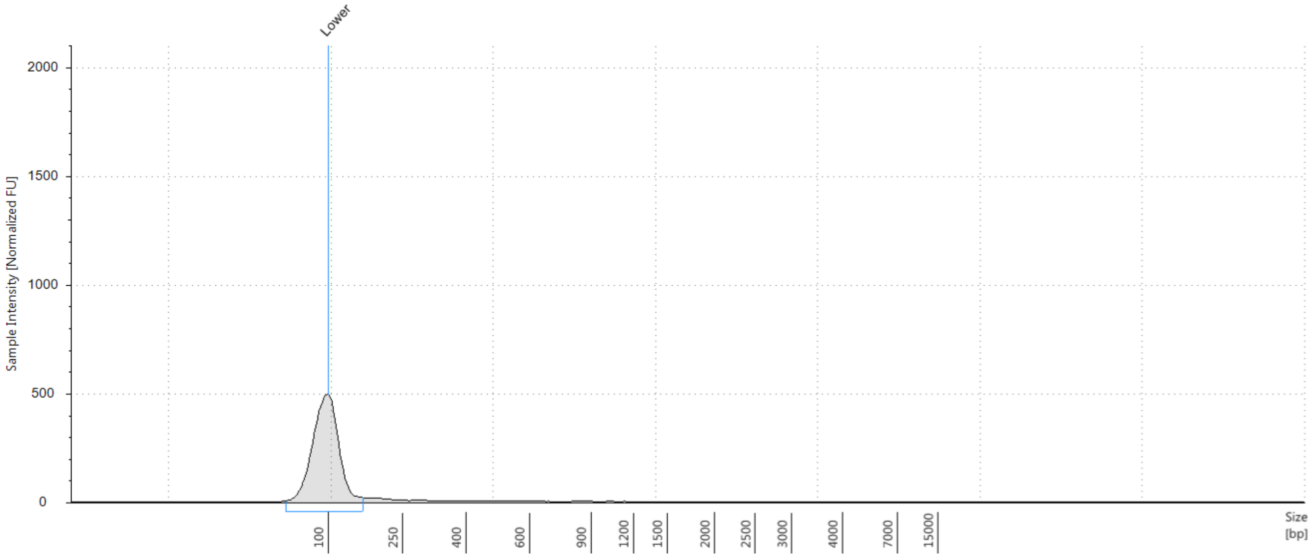


**Sample Table**

| Well | DIN | Conc. [ng/µl] | Sample Description | Alert | Observations |
|---|---|---|---|---|---|
| H1 | - | 1.46 | EB buffer | ⚠ | Issue with ladder peak detection (too few peaks detected); Caution! Expired ScreenTape device; Sample concentration outside functional range for DIN |

**Peak Table**

| Size [bp] | Calibrated Conc. [ng/µl] | Assigned Conc. [ng/µl] | % Integrated Area | From [bp] | To [bp] | Peak Comment |
|---|---|---|---|---|---|---|
| 100 | 8.50 | 8.50 | - | 14 | 168 | |

 © Agilent Technologies, Inc. 2017 
